# Cell type specific labeling and partial connectomes of dopaminergic circuits reveal non-synaptic communication and large-scale axonal remodeling after exposure to cocaine

**DOI:** 10.1101/2020.09.29.318881

**Authors:** G.A. Wildenberg, A.M. Sorokina, J.L. Koranda, A. Monical, C. Heer, M.E. Sheffield, X. Zhuang, DS McGehee, N. Kasthuri

## Abstract

Dopaminergic (DA) neurons exert profound influences on behavior including addiction. However, how DA axons communicate with target neurons and how those communications change, for example, with drug exposure, remains poorly understood. We combine recent advances in cell type specific labeling for electron microscopy with large volume three-dimensional serial electron microscopy – ‘connectomics’- to detail DA connections in the Nucleus Accumbens (NAc) across multiple animals and after exposure to cocaine. We find that DA axonal varicosities are of four general types: 38% are empty, 25% contain few small (∼50 nm diameter) vesicles, 19% contain few large (∼133 nm diameter) vesicles, and 18% have mixed small and large vesicles, suggesting that DA axons may use multiple types of neurotransmitters. Individual DA axons were significantly more likely to contain multiple varicosities of the same type relative to chance, suggesting a new method of DA axon classification. Across all categories, we find only rare examples (<2%, 6/410) of varicosities making specific synapses with any neighboring neuron with the few examples being made exclusively on the shafts and soma of resident NAc neurons. Instead, we find much more frequently (15%) that DA varicosities form spinule-like structures: physical membrane interdigitations with nearby dendrites or excitatory and inhibitory axons. Days after a brief exposure to cocaine, DA axons were extensively branched relative to controls but with similar densities of varicosities and spinules. Additionally, cocaine exposure results in the formation of blind-ended “bulbs” in DA axons, filled with mitochondria, and reminiscent of axonal retraction in the developing and damaged brain. Every bulb was surrounded by elaborated glia further suggestive of active remodeling. Finally, mitochondrial lengths increased by ∼2.2 times relative to control throughout DA axons and NAc spiny dendrites after cocaine exposure but not in DA soma or DA dendrites. We conclude that DA axonal transmission is unlikely to be mediated via classical synapses in the NAc and that the major locus of anatomical plasticity of DA circuits after exposure to cocaine are large scale axonal rearrangements with correlated changes in mitochondria.

## Introduction

The dopaminergic system, like many neuromodulatory systems of the brain, profoundly influences behaviors including learning risks and rewards, social cooperation, goal directed behavior, and decision making (Alcaro et al., 2007; Eskenazi et al., 2021; Liu et al., 2021). Moreover, alterations in dopaminergic circuitry in in animal models and humans are a hallmark of addictive behaviors and are likely the result of changes in how DA neurons communicate with targets (Berke and Hyman, 2000). However, while much is known about the molecular composition and functional roles of DA neurons in normal behavior and addiction (Alcaro *et al*., 2007; Beeler et al., 2009; Berke and Hyman, 2000; Liu and Kaeser, 2019; Liu et al., 2018), much less is known about how DA neurons physically connect with targets and how those ‘connections’ change with exposure to drugs of abuse.

There were several problems preventing progress. One is that, until recently, collecting and analyzing large volumes of brain tissue with electron microscopy, the gold standard for delineating neuronal connections, had been difficult and laborious. Recent advances in automation of the data collection (Hayworth et al., 2020; Kasthuri et al., 2015a; Xu et al., 2017; Yin et al., 2019) and algorithms to analyze the resulting terabytes of data have made such approaches more accessible (Funke et al., 2019; Januszewski et al., 2018; Saalfeld et al., 2012; Turaga et al., 2010). A second problem is that, unlike ionotropic excitatory or inhibitory connections, there is much less known about what potential DA synapses appear like at the ultra-structural level, where they occur on neurons, or how frequently (Liu *et al*., 2018; Rice and Cragg, 2008). Thus, analyzing DA circuits has required specific labeling, either with immunohistochemistry (Bérubé-Carrière et al., 2012; Moss and Bolam, 2008; Omelchenko and Sesack, 2009) (Liu *et al*., 2018), or genetic targeting (Dos Santos et al., 2018; Melchior et al., 2021; Mingote et al., 2019; Nasirova et al., 2021; Poulin et al., 2018). However, fundamental questions about neuromodulatory circuits remain unanswered: 1. how often do DA axons make synapses that appear like classic chemical synapses (i.e. with synaptic vesicles and post-synaptic densities), 2. where do DA synapses occur on target neurons, 3. what do DA axonal varicosities contain, and 4. are there signs of other physical interactions between DA axons and their targets?

Connectomics, the large scale and comprehensive 3D reconstructions of neurons and their connections, has emerged as a useful tool for understanding neural circuitry, revealing insights about how neurons connect that could not have been achieved any other way (Bae et al., 2018; Bates et al., 2020; Briggman et al., 2011; Helmstaedter et al., 2013; Karimi et al., 2020; Kasthuri *et al*., 2015a; Morgan and Lichtman, 2020). However, most current connectomic efforts in mammals center on cortex and focus almost exclusively on glutamatergic and GABAergic circuits (Gour et al., 2020; Karimi *et al*., 2020; Kasthuri *et al*., 2015a; Motta et al., 2019; Shapson-Coe et al., 2021). A potential solution would be to label DA axons in large volume EM datasets and recent advances in protein engineering have created such genetic labels (Clarke and Royle, 2018; Shu et al., 2011). One label, based on an *ascorbate peroxidase* from plants, Apex2 (Martell et al., 2017), has been expressed in targeted neuronal populations (Joesch et al., 2016; Martell et al., 2017; Sampathkumar et al., 2021; Tsang et al., 2018) and used to reconstruct connections of specific neurons millimeters away from their soma (Sampathkumar et al., 2021). But such approaches have not been applied to neuromodulatory circuits.

Here, we combine these new approaches to ask whether connectomes containing labeled DA circuits could reveal heretofore unknown details about how DA axons connect with targets. We labeled VTA dopamine neurons in 2 cocaine sensitized and 3 control mice using the genetically encoded pea peroxidase gene, Apex2, and created serial EM datasets of dopamine neurons in the VTA and their axons in the NAc (∼0.5mm x 0.5mm x 0.03mm volumes for each brain region) using the ATUM approach (Kasthuri *et al*., 2015a). Moreover, since DA pathways appear highly plastic (Calabresi et al., 2007; Jay, 2003; Pignatelli and Bonci, 2015; Pignatelli et al., 2017), especially with exposure or addiction to drugs of abuse (Berke and Hyman, 2000; Lüscher, 2016; Nestler, 2001; Saal et al., 2003; Ungless et al., 2001), we wondered whether connectomics could reveal potential structural changes of that plasticity. We focused on potential early changes after brief exposure to cocaine, testing the sensitivity of large volume EM for revealing acute changes. Furthermore, because EM datasets visualize every cell membrane and sub-cellular organelle, we hypothesized that potential structural changes in DA axons could be correlated with potential structural changes in other cell types and subcellular organelles (i.e., glia, mitochondria, or target neurons in the same EM datasets). The ability to discover such correlations are difficult, if not impossible, in either small volume EM stacks or fluorescently labeled optical datasets and therefore would demonstrate the value of comprehensive connectomes for the study of structural correlates of neuromodulatory plasticity.

We found several novel aspects of DA axonal biology. First, most (∼98%) DA axonal varicosities showed no clear ultrastructural signs of specific synapses with target neurons (i.e., PSDs adjacent to vesicle clouds, parallel membranes, etc.). Rather, DA axons generally contained four types of varicosities: empty, small vesicles, large vesicles or mixed (i.e. small and large), and individual axons were more likely to contain multiple examples of a specific varicosity type than expected by chance. The rare evidence of an ultrastructural synapse occurred exclusively on the shaft and soma of resident NAc neurons. Additionally, far more common than ultrastructural signs of specific synapses were ‘spinules’, where the membranes of DA axonal varicosities interdigitated with the membranes of nearby excitatory and inhibitory axons as well as dendrites of resident neurons, reminiscent of spinules in other parts of the brain (Petralia et al., 2015; Spacek and Harris, 2004; Tao-Cheng et al., 2009; Tarrant and Routtenberg, 1977; WESTRUM and BLACKSTAD, 1962).

Brief cocaine exposure caused widespread branching of DA axons with new branches containing similar rates of varicosities and spinules. Surprisingly, ∼ 50% of DA axons contained branches ending in blind ‘bulbs’, reminiscent of retraction bulbs in the developing and injured brain (Balice-Gordon et al., 1993; Bishop et al., 2004; Bixby, 1981; Canty et al., 2020; Ertürk et al., 2007; Johnson et al., 2013; Korneliussen and Jansen, 1976; Riley, 1981), and often surrounded by elaborated processes of nearby glia with signs of axonal engulfment. Also, axonal changes were accompanied by targeted changes in mitochondria morphology: mitochondria in DA axons and their putative targets were 2.2-fold longer but we found no change in mitochondrial length in NAc glutamatergic afferent axons, nor in VTA DA soma or DA dendrites. Overall, our findings demonstrate advantages to using connectomic tools to give new insights into DA axon biology and how brief exposure to cocaine impacts mesoaccumbens dopamine circuitry.

## Results

To selectively label mesoaccumbens dopamine neurons, we bilaterally injected Adeno-associated virus (AAV) encoding either a CRE-dependent cytosolic or mitochondrial targeted pea peroxidase, Apex2 (Martell *et al*., 2017; Sampathkumar *et al*., 2021), into the VTA of 5 mice expressing CRE recombinase from the promoter of the dopamine transporter gene (DAT-CRE) (Backman et al., 2006) (**Figure 1A**). Four weeks following AAV expression, mice were perfused, and brain slices were treated with 3,3′-Diaminobenzidine (DAB) to convert Apex2 into a visible precipitate with binding affinity to osmium tetroxide (**Figure 1B**) (Joesch *et al*., 2016; Martell *et al*., 2017). Brain sections with DAB precipitate localized to the appropriate brain regions were cut into smaller pieces surrounding the VTA and the medial shell of the NAc (**Figure 1B**, green boxes) and further processed for serial EM (Hua et al., 2015) (see Methods). We focused our analysis on DA axons projecting to the medial shell of the NAc (referred to as simply, NAc) because: 1. It receives the majority of VTA projecting DA axons (Beckstead et al., 1979), 2. It is reported to undergo functional and structural plasticity in response to cocaine (MacAskill et al., 2014), and 3. areal landmarks allow for unambiguous identification of NAc across replicates (**Figure 1B**, example landmarks depicted with red ovals highlighting negative staining of the anterior commissure and nuclei along midline of the brain, and see **Methods**). We first collected single 2D EM sections to confirm that both cytosolic- and mitochondrial-localized Apex2 unambiguously labeled DA neurons at their soma (**Figure 1C-D**, left panels, red arrow), dendrites (**Figure 1C-D**, middle panels, cyan arrow) and small axonal processes several millimeters away from the injection site (**Figure 1C-D**, right panel, yellow arrow). We found no evidence of labeled soma in any other brain region, including the NAc, suggesting that the viral strategy worked, targeting only genetically specified DA axons in an anterograde manner. After confirming appropriate Apex2 expression for all mice used in this study, ∼2000, 40nm thick sections were collected from both the VTA and NAc, and volumes of ∼0.5mm x 0.5 mm x 0.03mm were imaged by serial electron microscopy.

**Figure 1.**
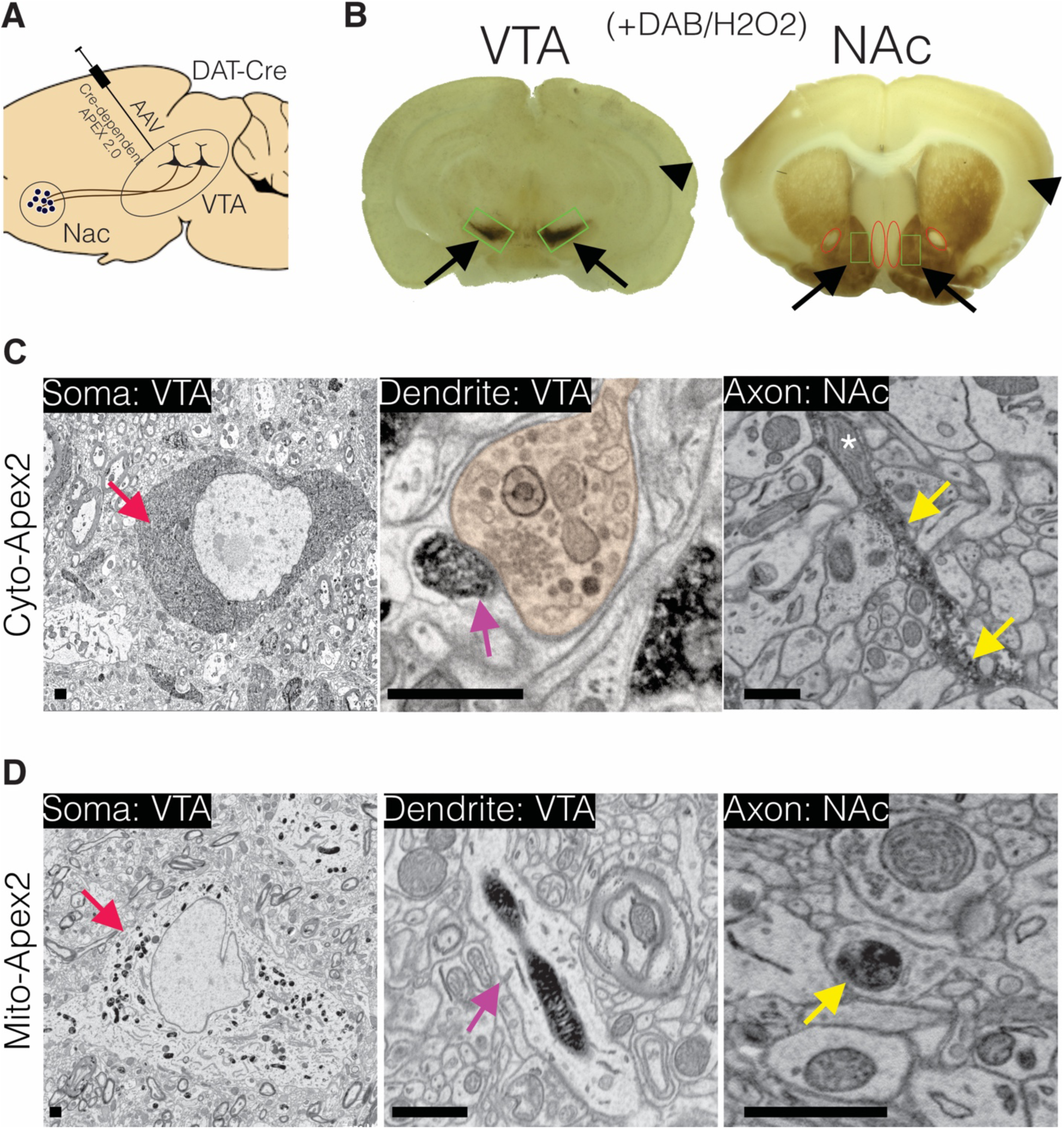
Experimental design for Dopamine connectomics. (**A**) AAVs expressing Cre-dependent Apex2 were bilaterally injected into the Ventral Tegmental Area (VTA) of transgenic mice expressing Cre in Dopamine Transporter positive neurons (DAT-CRE). (**B**) ∼4 weeks after AAV injections, vibratome sections (∼300 microns thick) show strong Apex2 labeling in VTA and NAc after staining with 3’3’-Diaminobenzidene (DAB) and Hydrogen Peroxide (H202) before EM processing (see Methods). Black arrows point to an Apex2 positive region and black arrowhead points to an Apex2 negative region. Green rectangles highlight the VTA and medial shell of the NAc region dissected out and processed for EM. Red ovals highlight areal landmarks to ensure the same region was dissected across all animals. (**C-D**) Representative EM images of cytoplasmic (C, top row) and mitochondrial (D, bottom row) Apex2+ DA neurons (Left panel: Apex2 Soma in the VTA (red arrows); *Middle panel*: Apex2 top panel shows a DA dendritic spine forming a synapse (purple arrow) with presynaptic bouton (orange) in the VTA, and the bottom panel shows a narrow DA dendrite expressing mitochondrial Apex; *Right panel*: Apex2 axon in the NAc with narrow (yellow arrow) and thick varicosities (yellow arrowhead). Cytosolic Apex2 does not obscure mitochondria (top panel, asterisk), and mitochondrial Apex2 (bottom) only fills up mitochondria. Scale bar = (C,D) 1 μm.

We first described ultra-structural features of putative DA contacts with target neurons (i.e. what, if any, types of synapses do DA axons make?). We characterized DA axonal varicosities (see Methods) as the most likely location along axons for evidence of synaptic contacts in DA axons in ‘control’ mice not subject to any of the behavioral experiments used in the cocaine sensitization experiments described below. We used mito-Apex2 to clearly visualize ultrastructural features (e.g. vesicle clouds, PSD), which might be obscured by cytoplasmic Apex2 expression. We reconstructed 410 varicosities on 75 individual DA axonal fragments in one mouse (p105, male). We defined varicosities as regions where the DA axon diameter increased ∼3 fold relative to the thinnest portion of the axon, either in the absence of mitochondria, or in the presence of a mitochondria when accompanied by vesicles within the same varicosity. Broadly, varicosities were of four types based on their contents: 1. empty (38%: 156/410), 2. several small vesicles (25%: 101/410; 48 ± 1.7 nm in diameter), 3. few, large vesicles (19%: 78/410; 133 ± 5.5 nm in diameter), or 4. mixture of large and small vesicles (18%: 75/410) (**Figure 2A**). **Figure 2B** shows representative examples of each varicosity type on a single axon colored to correspond to the pie chart of all varicosity distributions in **Figure 2A** (red arrows or arrowheads point to large and small vesicles, respectively, white arrow pointing to an empty varicosity). Varicosities usually contained a small number of vesicles (∼15-33 small vesicles/∼5-11 large vesicles) as compared to nearby putative cortical synapses innervating nearby spines (150-200 vesicles). Although individual axons often contained more than one type of varicosity (**Figure 2C**), across many axons, we found axons with long stretches of a particular varicosity type (**Figure 3A**), suggesting a preference of individual axons for a specific type of varicosity. Thus, we asked whether such distributions along individual axons were consistent with random sampling of varicosity type, weighted by their population frequencies (i.e. Figure 2A). Thus, we performed a Monte Carlo simulation (Rubinstein and Kroese, 2008) such that, for every DA axon with three or more varicosities reconstructed from the real dataset, a DA axon was simulated with the exact number of varicosities/axon so that all axons in the real data set had an exact replicate in the simulation. For every trial of the simulation (100,000 trials), we randomly assigned a varicosity type, based on their population frequency, to each varicosity of a simulated DA axon. Finally, we asked how often in the 100,000 trials do simulated axons have long stretches of a particular varicosity (i.e. how many simulated axons have more than 1, more than 2, or more than 3, etc. of Type I, II, III, or IV varicosity). We found that actual DA axons were far more likely to contain long stretches of all varicosity types, relative to the simulated population (**Figure 3B**). For example, Type II, III, and IV varicosities appeared multiple times along single axons (e.g. >3 instances), which rarely occurred in over 100,000 trials of the simulation (see **Figure 3-Figure Supplement 1** for all P-values). We conclude that individual ‘real’ DA axons have multiple instances of the same varicosity type which are unlikely to have occurred by chance and suggest that DA axons can be classified by the types of varicosities present along an axon.

**Figure 2.**
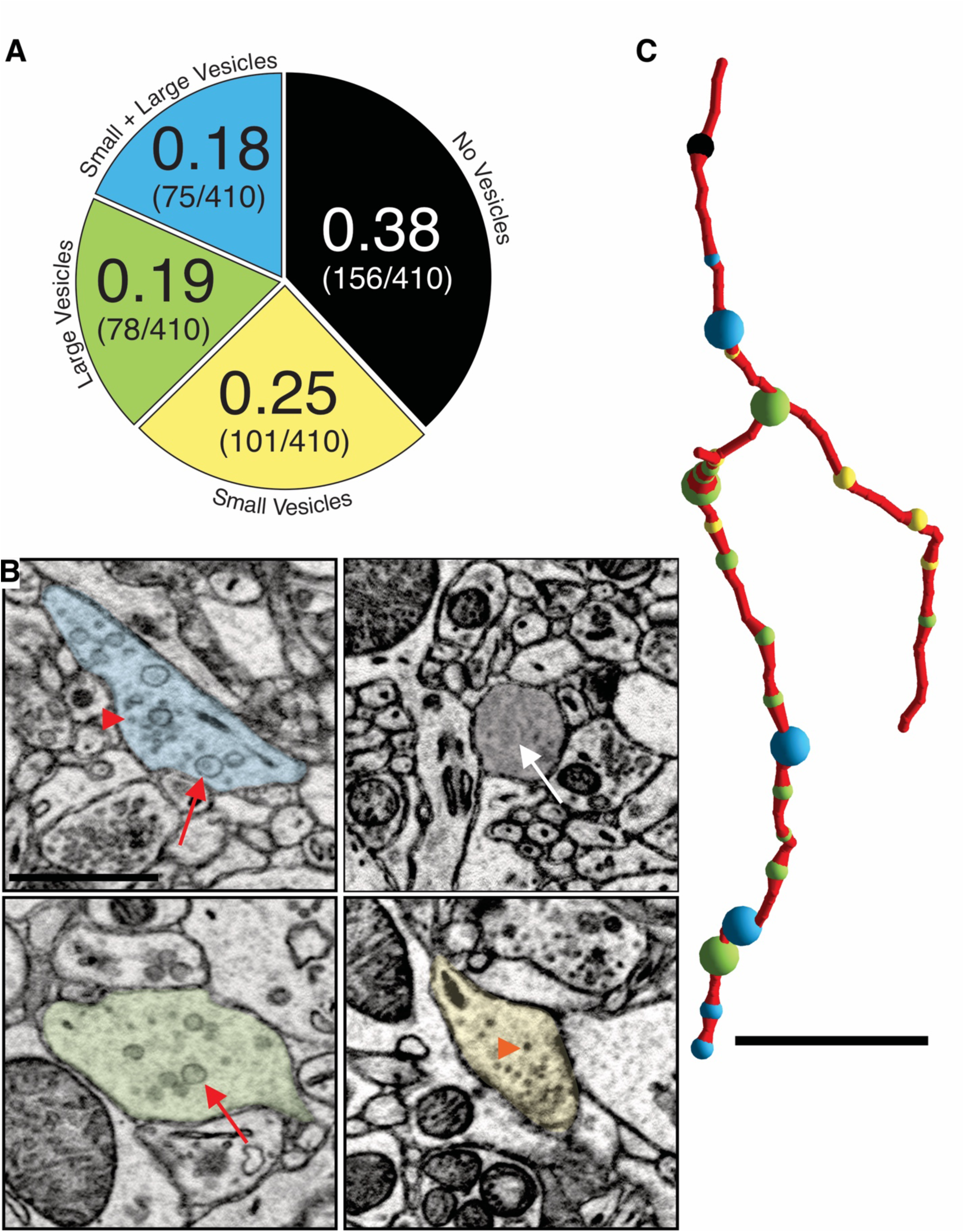
DA axon varicosities are either empty or filled with different sized vesicles. (**A**) Pie chart showing ratio of each kind of DA varicosity across a population of DA axons (n = 409 varicosities scored across 75 axons, 1 mouse). (**B**) Single 2D images showing representative examples of each kind of DA axon varicosity with color overlay corresponding to each type (refer to pie chart in A). Red arrowheads point to small vesicles, red arrows point to large vesicles, and the white arrow points to an empty varicosity. (**C**) Reconstruction of a single Mito-Apex2 DA axon with each class of varicosity marked as a differently colored spheres: black = no vesicles, yellow = small vesicles, green = large vesicles, and blue = small and large vesicle filled varicosity. Spheres are scaled to the size of the bouton. Scale bar (B) = 1 µm, (C) = 10 µm.

**Figure 3.**
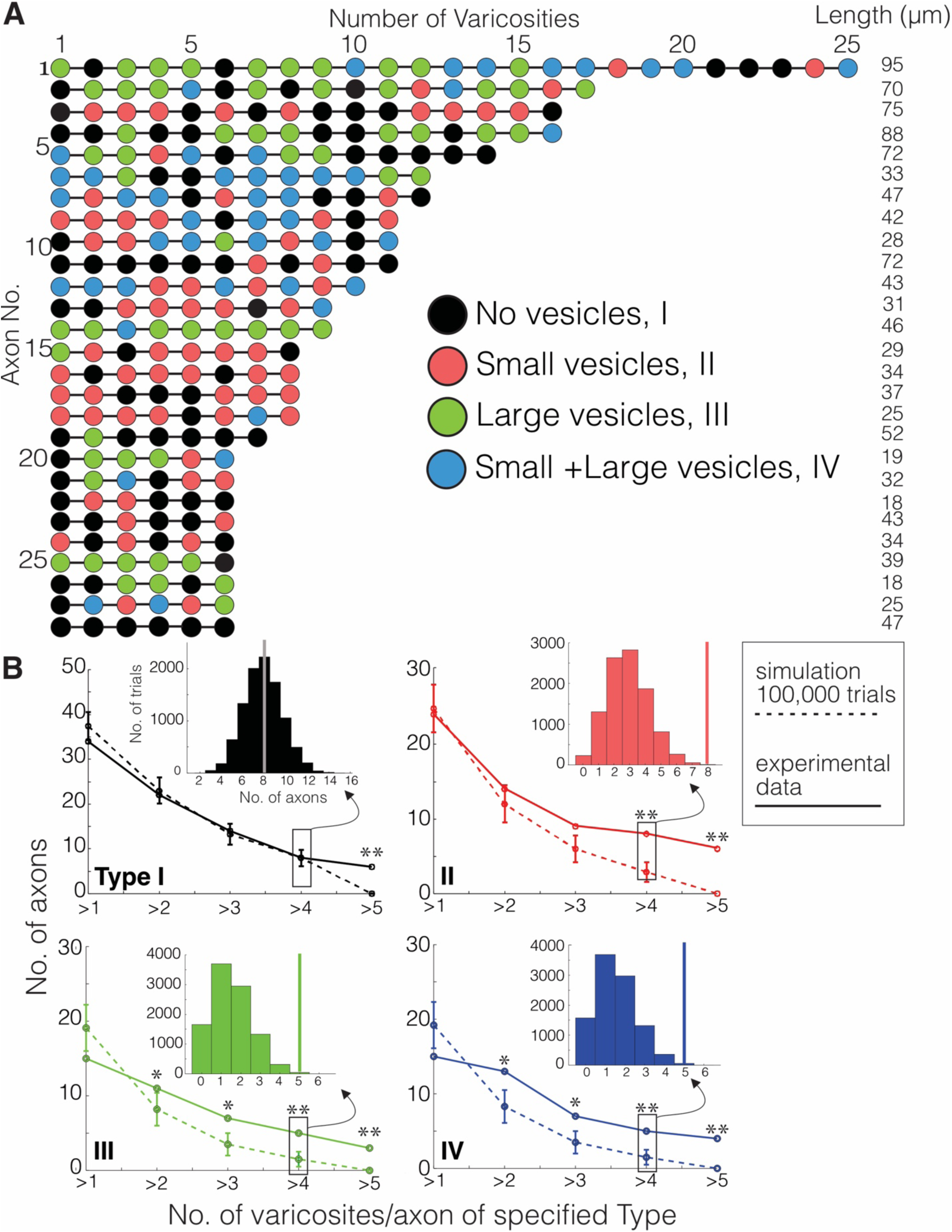
Monte Carlo simulation of DA varicosity types. (**A**) Mito-Apex2 DA axons containing 6 or more varicosities in the field of view are depicted with the linear order of their varicosities. Each varicosity is shown as a colored circle with each color representing a different varicosity type. The length (µm) of each reconstructed DA axon is listed on the right. (**B**) For every DA axon with three or more varicosities reconstructed from the real dataset, a simulated DA axon was created to match the number of varicosities/axon (e.g. a simulated DA axon was made with 3 varicosities to match a real DA axon with 3 varicosities). A Monte Carlo simulation was then ran 100,000 times to randomly assign varicosity types based on their population frequency reported in Figure 2A. For both simulated (dashed line) and real (solid line) DA axons, the number of axons was plotted against the number of varicosities/axon containing the specified type for each graph (e.g. in the top left graph, all axons containing more than 1 Type I varicosity were counted at the “>1” position of the x-axis). *Inset*: example of the simulated distribution and real axon (solid vertical line) data generated for these analyses for the number of axons that had more than 4 varicosities of each type. Asterisks denote statistically significant differences between the simulated and real data when the p-value is between 0.01 and 0.05 (*) or lower than 0.01 (**). P-values are shown for each data point in Figure 3 Supplement 1.

Finally, given the lack of obvious synaptic structures, we asked whether the ∼62% of varicosities with vesicles, but no obvious PSDs, could serve as putative sites of volume transmission: where local DA release modulates synaptic transmission between, for example, ionotropic glutamatergic synapses of nearby cortical axons and dendritic spines of NAc medium spiny neuron (MSN) (Agnati et al., 1995; Liu *et al*., 2021; Rice and Cragg, 2008). Thus, we surveyed the immediate 3D space around DA varicosities and scored them for membrane-to-membrane apposition to either the presynaptic axonal terminal or postsynaptic spine/shaft of all types of synapses (e.g. soma, shaft, or spine). We found little evidence that vesicle filled varicosities were more likely to be physically adjacent to a synapse than an empty varicosity (+vesicle varicosities: 156/249 (63%); -vesicle varicosities: 104/156 (66%)), and additionally we found no evidence for DA varicosity making synapses with spines, reducing the possibility that DA axons target volume transmission to nearby spines.

In rare instances (6/410), we found clear ultrastructural evidence (i.e. clusters of vesicles with clear PSDs in the target neuron, parallel apposition of presynaptic axonal and target neuron membranes) of DA axons making synapses on the soma and dendritic shaft of resident NAc neurons. **Figure 4** shows three examples of mito-Apex2+ DA axons forming synapses on soma and the shaft of resident NAc neurons. In each case, the DA axons varicosity had a prominent vesicle cloud localized around a PSD (red arrows). These results demonstrate both that our approach can detect rare DA synapses and also that the majority of DA varicosities show no evidence of widespread structural connections with other neurons.

**Figure 4.**
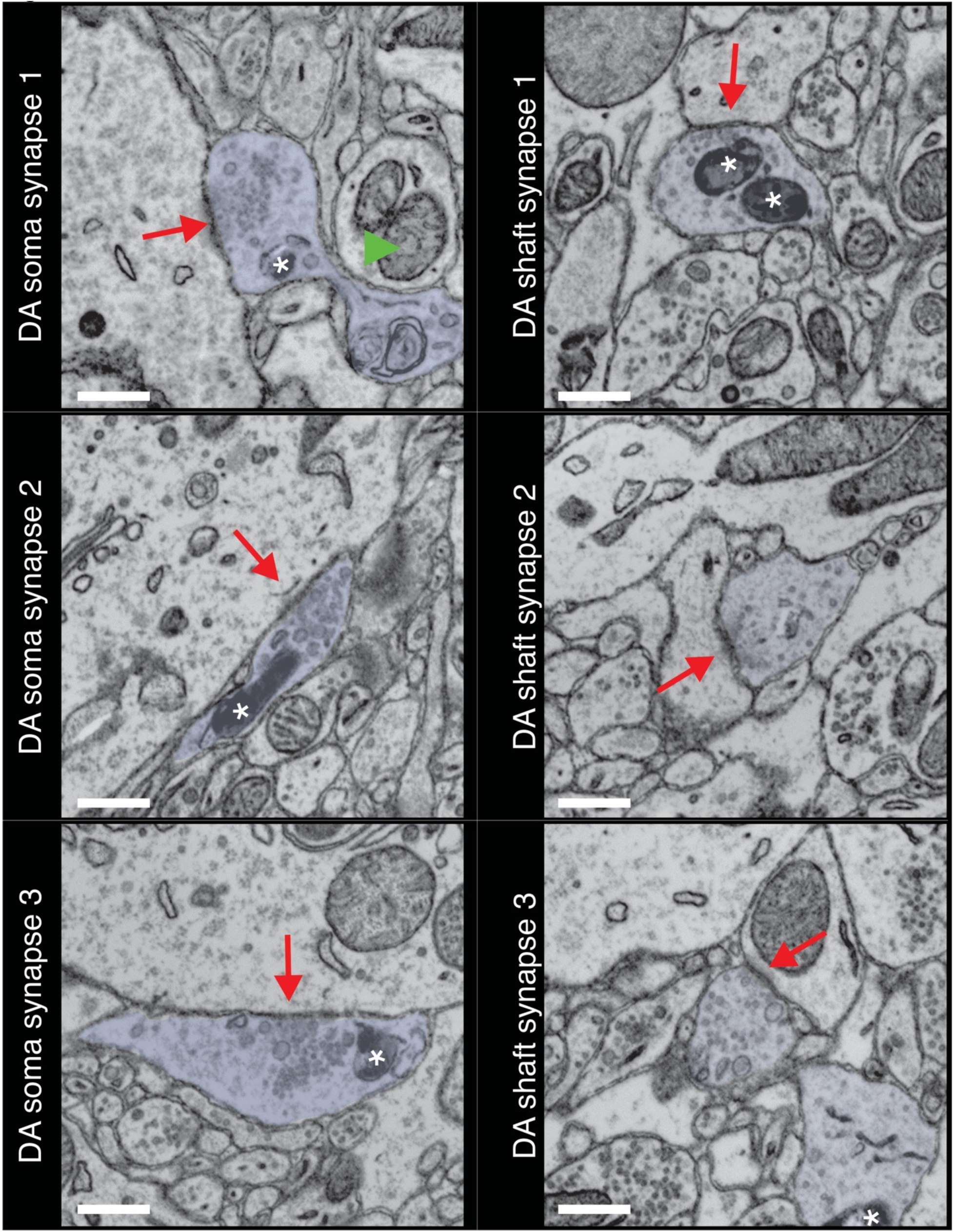
DA axons make synapses on the soma and shaft of NAc resident neurons. Three examples of DA axons making synapses on the soma (left column) and shaft (right column) are depicted. Mito-Apex2 DA axons are shaded in light blue with an asterisk marking an Apex2 positive mitochondria which are significantly darker than Apex2 negative mitochondria (green arrowhead). Red arrows point to the PSD formed between the DA axon and soma. Scale bar = 500 nm.

Lastly, we asked whether DA axons physical interact in any other way with other neurons in the volume. We found in mito-Apex2 expressing DA axons that 15% (61/410) of varicosities contained membrane invaginations with either unlabeled axons or dendrites. Representative examples are shown as image montages and 3D reconstructions from both mito- and cyto-Apex2 datasets (**Figure 5** and **Figure 5 – Figure Supplement 1**). Furthermore, these invaginations or “contact points” were made primarily between DA axons and unlabeled axons, most anatomically similar to chemical synapse (CS) forming axons (e.g. Glutamatergic or GABAergic axons) (83%, 49/59) and a smaller proportion with dendrites (17%, 10/59). The unlabeled CS axons that made invaginations into DA axons could be further classified into a cohort (∼43%) making additional synapses on dendritic spines, suggesting they were excitatory and the rest, 57%, made synapses on dendritic shafts with little sign of a PSD, suggesting they were inhibitory. All dendrites with contact points with DA axons were 100% spiny. These invaginations closely resemble previously characterized structures called spinules which are believed to be postsynaptic projections into presynaptic terminals (Petralia *et al*., 2015; Spacek and Harris, 2004; Tao-Cheng *et al*., 2009; Tarrant and Routtenberg, 1977; WESTRUM and BLACKSTAD, 1962) and have recently been shown to be induced by neuronal activity (Tao-Cheng *et al*., 2009; Zaccard et al., 2020) (see Discussion).

**Figure 5.**
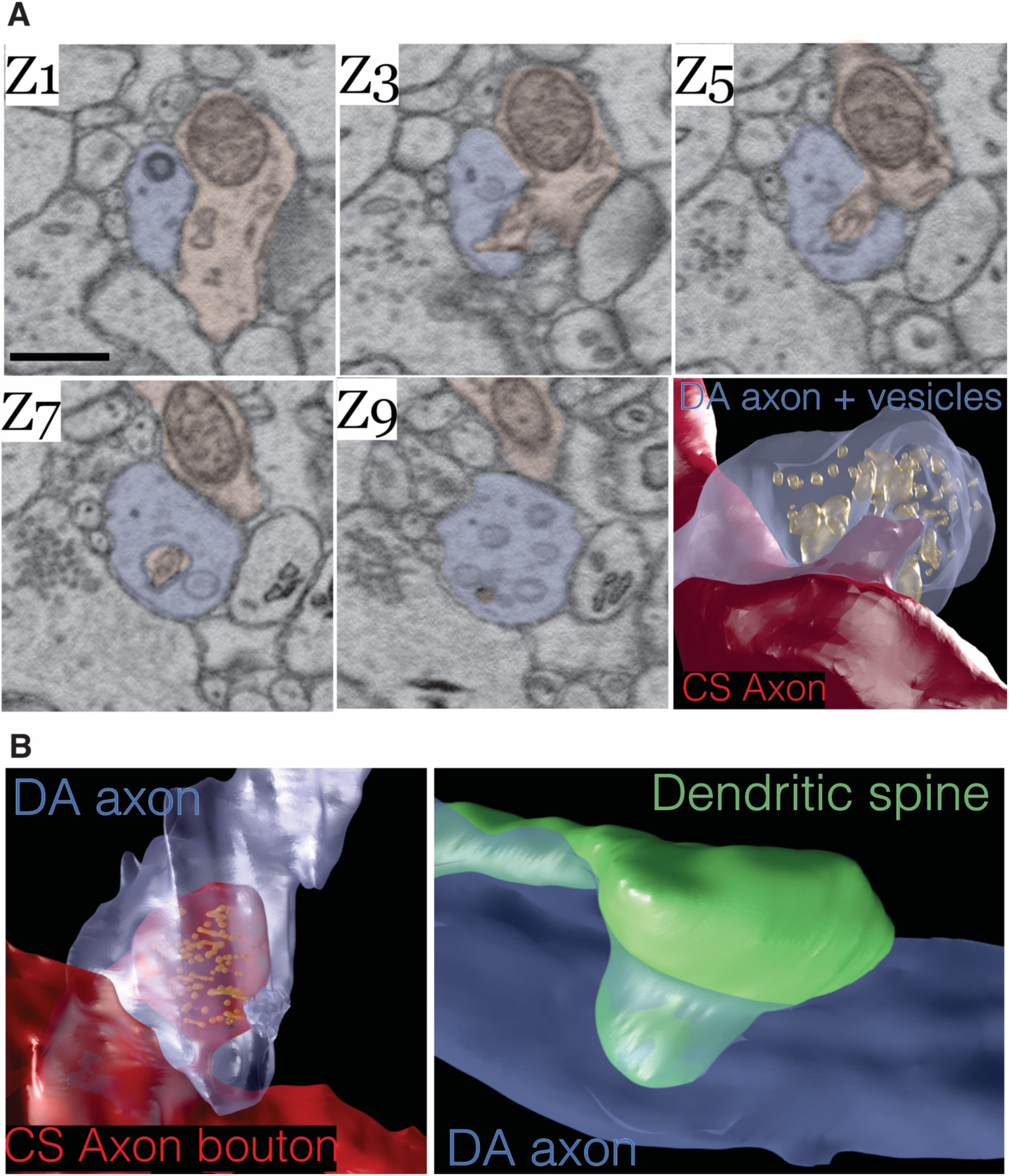
Dopamine contact points are physical interdigitations between DA axons and either afferent axons or NAc dendrites. **(A)** image montage of every other EM serial section (Z1-Z9) from a mouse expressing mito-Apex2 in DA neurons. Highlighted in blue is the mito-Apex2 DA axon, and in red is the interdigitating chemical synapse (CS) axon. *Bottom right*: 3D rendering showing interdigitation of a CS axon (red) into the vesicle filled DA axon (blue). (**B**) 3D renderings of two other examples of DA axon interdigitations. A vesicle filled CS axon (red) or dendritic spine (green) interdigitate into a DA axon (blue) in the left and right images, respectively. Scale bar = (A) 1 µm.

Given this basic appreciation of DA circuits, we next asked whether connectomic datasets could reveal changes in the physical circuitry of DA axons after exposure to cocaine. Two DAT-CRE mice expressing cytosolic Apex2 after AAV injections were subjected to a cocaine sensitization and associative conditioning protocol previously shown to induce morphological changes in the mesoaccumbens dopamine pathway (Beeler *et al*., 2009; Li et al., 2004; Singer et al., 2009) and compared to two controls injected with equivalent volumes of saline and exposed to the same associative conditioning protocol. Briefly, mice were given one daily intraperitoneal (IP) injection of either cocaine (10 mg/kg, n=2) or an equivalent volume of saline (n=2) every other day for a total of 4 injections (**Figure 6-Figure Supplement 1A**) and assessed for locomotor activity in a novel environment. Consistent with prior reports, cocaine treated mice showed an increase in daily activity relative to the control with each successive treatment (**Figure 6-Figure Supplement 1B, C**). After the final injection, mice went through a standard four-day abstinence period, before being processed for Apex2 staining and large volume serial EM. A four-day abstinence period was chosen to minimize potential acute effects of cocaine on dopamine circuitry, focusing this study on potentially more long-term changes caused by cocaine sensitization (Benuck et al., 1987; Eipper-Mains et al., 2013; Grimm et al., 2001; Hollander and Carelli, 2007; Nestler, 2001; Parsons et al., 1991).

We first analyzed the trajectories of axons in 20 nm (x, y) resolution EM volumes from NAc datasets of 2 control (+saline) and 2 cocaine treated (+cocaine) mice expressing cyto-Apex2. cyto-Apex2 was used for this analysis because it allowed us to follow thin axons at lower resolutions. We traced 85 cyto-Apex2+ DA axons (44 from controls, 41 from cocaine treated, total length of 5,192.8 microns) and immediately found that DA axons were highly branched in cocaine treated mice relative to controls (**Figure 6A**; +saline = blue axons, +cocaine = red axons). Across labeled DA axons, there was an average 4-fold increase in branch number for similar axon lengths with exposure to cocaine (**Figure 6B**; mean ± SEM branch number/µm length of axon: +saline, 0.01 ± 0.002, n = 44 axons, 2 mice; +cocaine, 0.04 ± 0.005, n = 41 axons, 2 mice. P = 3.96e-8). This increased branching was accompanied by an increased number of invaginating contact points (i.e. “spinules”) and varicosities in cocaine exposed animals as described above in Figure 2 and 5, but the number of contacts per length was similar. (**Figure 6C**; mean ± SEM number of contacts points (c.p.)/µm length of axon: +saline, 0.2 ± 0.02, n = 125 contact points across 10 axons, 2 mice; +cocaine, 0.2 ± 0.03, n = 142 contact points across 9 axons, 2 mice. P = 0.90). Interestingly, control axons showed regular spacing of such contacts along the axon (Figure 5A, left), whereas in cocaine exposed and highly branched DA axons, there were large regions of axon that were completely devoid of any contacts (Figure 5A, right). Finally, we did not see an obvious difference in which kind of neuronal process formed invaginating contact points (e.g., axo-axonic versus axo-dendritic) (**Figure 6-Figure Supplement 2**: control: 83% (49/59) axo-axonic, 17% (10/59) axo-dendritic. n = 59 contact points across 19 DA axons; cocaine: 77% (56/73) axo-axonic, 23% (17/73) axo-dendritic, n = 73 contact points across 20 DA axons). These results suggest that while there were large changes in axonal structure and the numbers of contact points, the nature of individual varicosities and interactions with target neurons remained similar.

**Figure 6.**
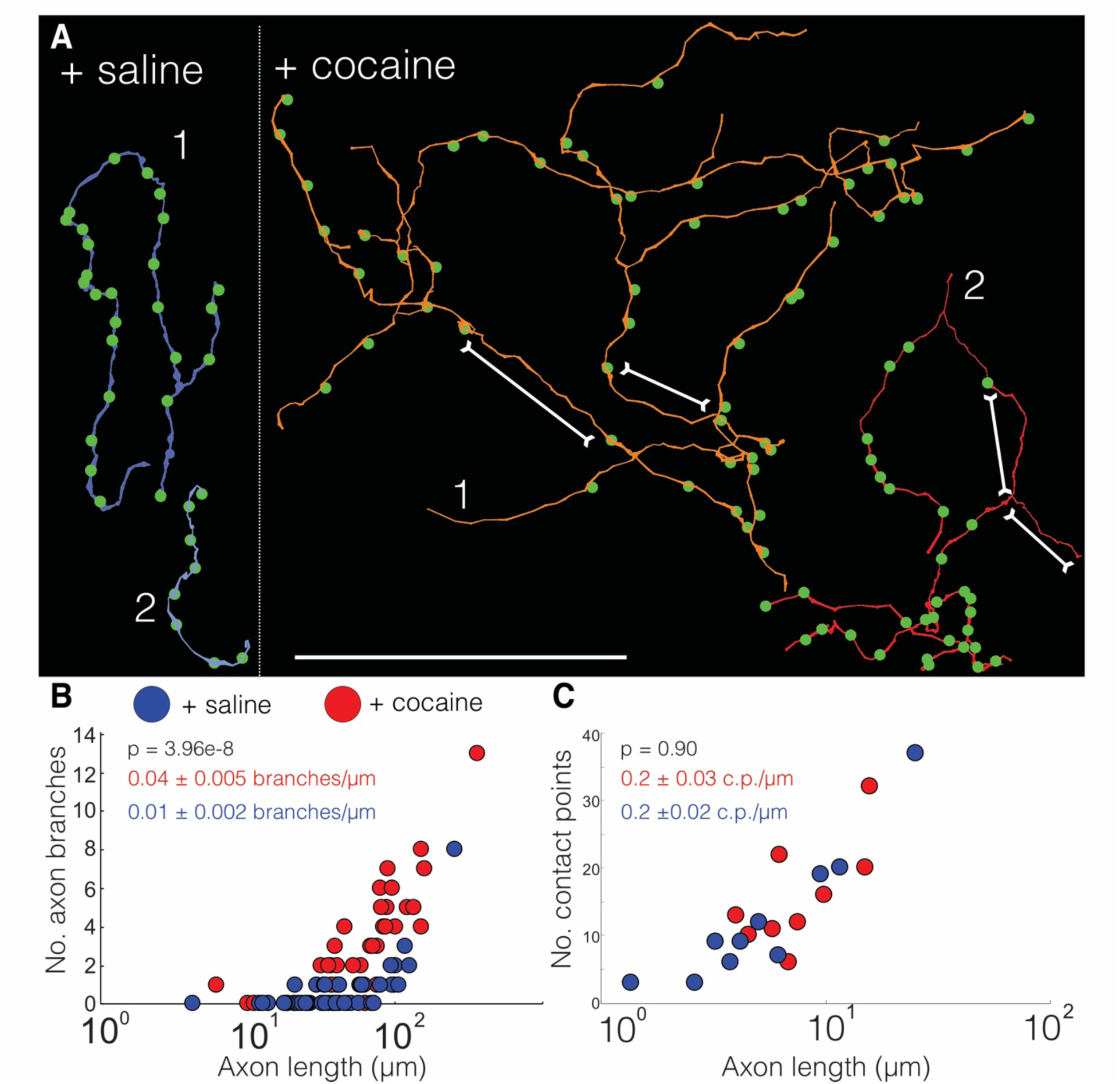
Cocaine increases branching of Apex2 dopamine axons in cocaine sensitized mice. **(A)** Two representative reconstructions of DA axons each from saline (left, blue) and cocaine (right, red) treated mice. Green circles represent contact points (i.e. spinules) where other neurons interdigitate with the DA axon. (**B**) Scatter plot of the number of DA axons branches versus axon length (μm) (+saline: 0.01 ± 0.002 branches/µm length of axon, n = 44 axons, 2 mice; +cocaine: 0.04 ± 0.005 branches/µm length of axon, n = 41 axons, 2 mice. P = 3.96e-8. (**C**) Scatter plot of the number of contact points (i.e. “spinules”) versus axon length (µm) (+saline: 0.2 ± 0.02 contact points (c.p.)/µm length of axon, n = 125 contact points over 10 axons, 2 mice; +cocaine: 0.2 ± 0.03 c.p./µm length of axon, n = 142 contact points over 9 axons, 2 mice. P =0.90). Data: mean ± SEM. *P*-values: two-tailed Mann-Whitney U test. Scale bar = (A) 40 µm.

The second obvious feature of DA axons exposed to cocaine was the occurrence of large axonal swellings or bulbs (**Figure 7**). The swellings were large (mean ± SEM diameter: 2.2 ± 0.3 µm, n = 23), significantly larger than varicosities in control animals (mean ± SEM diameter: 0.4 ± 0.02 µm, n = 118 varicosities) and at times reaching the size of neuronal soma (**Figure 7A**). These ‘bulbs’ were common in axons (∼56%, 17/30 axons) in two cocaine exposed animals and we did not see a single example in DA axons from two control animals (0/29 axons), suggesting that Apex2 expression alone does not cause these swellings (**Figure 7B**; mean ± SEM swellings/µm length of axon: + saline, 0.00 ± 0.0, n = 29 axons, 2 mice; +cocaine, 0.04 ± 0.02, n = 30 axons, 2 mice. P = 1.7e-5). **Figure 7C** shows reconstructions of two of these axons with swellings (large spheres with asterisk). DA axons were found to have either a large swelling along the axon (bottom reconstruction) often surrounded by medium sized swellings (green spheres), or contained terminal bulbs (top reconstruction, asterisk) reminiscent of axon retraction bulbs observed in developing neuromuscular junctions and damaged brains. Additionally, we did not observe any swellings in VTA DA dendrites despite also being sites of dopamine release (Liu and Kaeser, 2019) nor in any NAc dendrites or afferent axons (i.e. excitatory axons that make chemical synapses from cortical and subcortical areas) (data not shown). We next re-imaged EM volumes around different swellings at a higher resolution (∼6nm x, y) to better resolve the contents of these structures. The left image in **Figure 7D** shows a 2D EM image of a representative DA axon bulb (red) filled with mitochondria (2 examples highlighted in blue). When this bulb was reconstructed into a 3D rendering from the serial EM sections spanning the region, we found the mitochondria to be extremely elongated, twisted, and packed into the swelling (**Figure 7D**, *right*: the bottom half of the outer membrane rendering of the bulb is removed to visualize the internal mitochondria) and that nearly all bulbs examined were similarly packed with mitochondria (data not shown).

**Figure 7.**
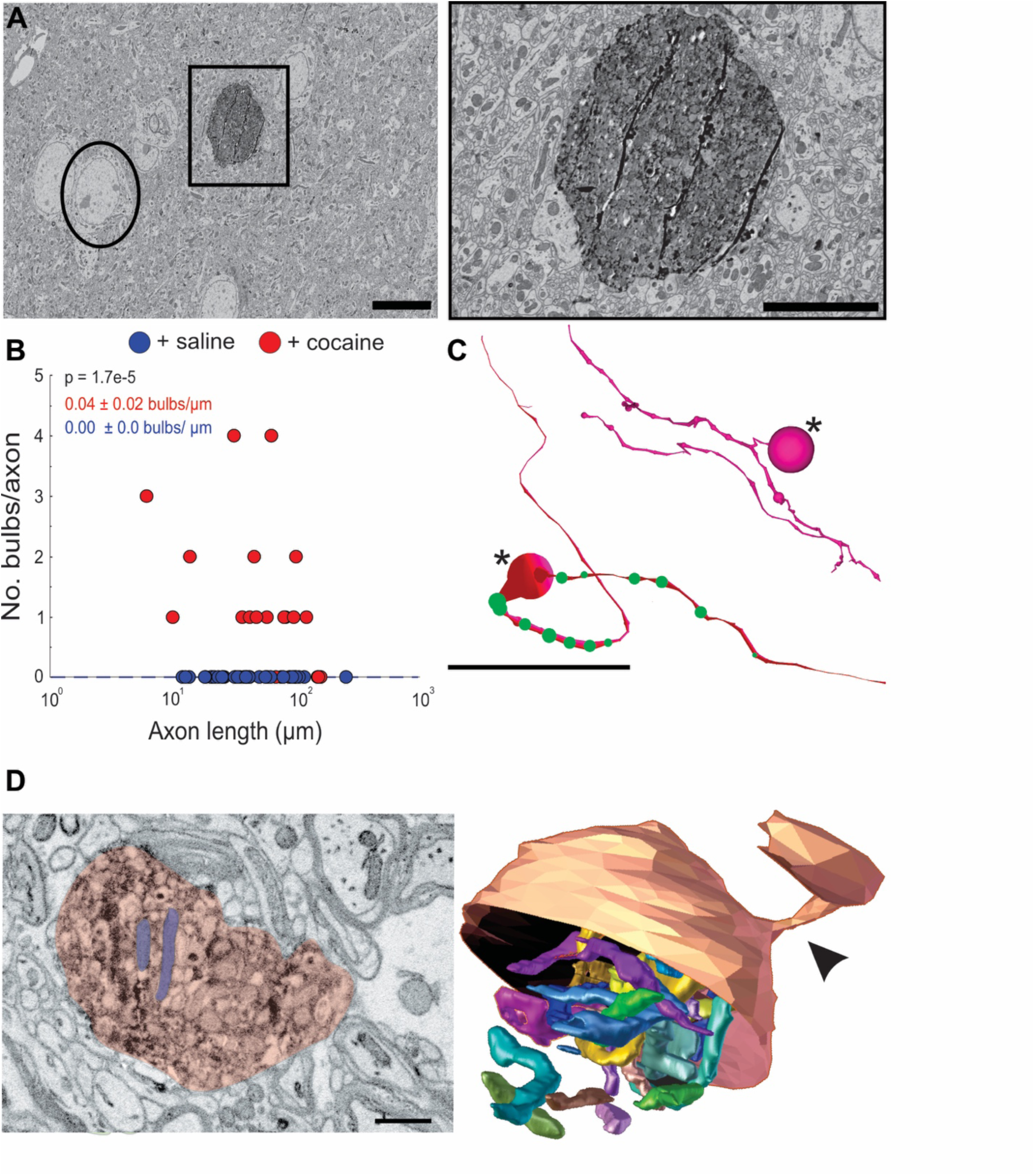
Cocaine results in the formation of large swellings filled with mitochondria in Apex2+ DA axons. (**A**) *Left*: 2D EM image of Apex2 labeled DA axon bulb in cocaine treated mouse (square) as compared to a neighboring neuronal soma (circle). *Right*: zoomed in 2D EM image of Apex2 DA axon bulb from left panel. (**B**) Scatter plot of the number of large swellings versus axon length (μm) (+ saline: 0.00 ± 0.0 swellings/µm length of axon, n = 29 axons; 2 mice + cocaine: 0.04 ± 0.02 swellings/µm length of axon, n = 30 axons, 2 mice. P = 1.7e-5. (**C**) Reconstructions of two representative Apex+ Dopamine axons with large swellings (asterisk) and medium sized swellings (green spheres). Top reconstruction depicts an axon with a terminal axon bulb and bottom reconstruction shows one along the axon. (**D**) *Left*: 2D EM image of Apex+ large DA axon swellings (red) filled with mitochondria (2 examples highlighted in purple) found in the NAc of cocaine treated animals. Both swellings are filled with mitochondria (examples highlighted in blue). *Right*: 3D segmentation of swelling and extremely long and coiled mitochondria found inside. Only the top half of the DA axon swelling is depicted to illustrate the mitochondria contained within. In this example, the swelling is at the end of the DA axon where it is attached to a thinner portion of the axon (arrowheads). Scale bar: (A) left: 10µm, right: 5 µm, (C) 20 μm, (D) 500 nm. Data: mean ± SEM. *P*-values: two-tailed Mann-Whitney U test.

Large axon bulbs have been most commonly associated with axonal pruning events where terminal boutons that lose their synaptic connection with a postsynaptic target form into large swellings (i.e. bulbs) and are engulfed by Schwann cells and other glial cell types presumably to remove the orphaned bouton (Bishop *et al*., 2004; Wilton et al., 2019). We also saw evidence of glial involvement surrounding DA axonal bulbs. **Figure 8A-B** shows a 3D reconstruction matched with a montage of EM images in **Figure 8C** of a representative DA axon bulb surrounded by numerous glial cells. Each glial cell was traced out in the volume to confirm they had characteristic non-neuronal, glial morphology (i.e. extensive branching, granules, etc., see Methods) (Fernández-Arjona et al., 2017; Heindl et al., 2018; Reichenbach et al., 2010), and that each traced object was an individual glial cell (i.e. the reconstructed processes came from different cells). Presence of several glia around the DA axon swelling provides further evidence that these are sites of active remodeling and plasticity in response to cocaine.

**Figure 8.**
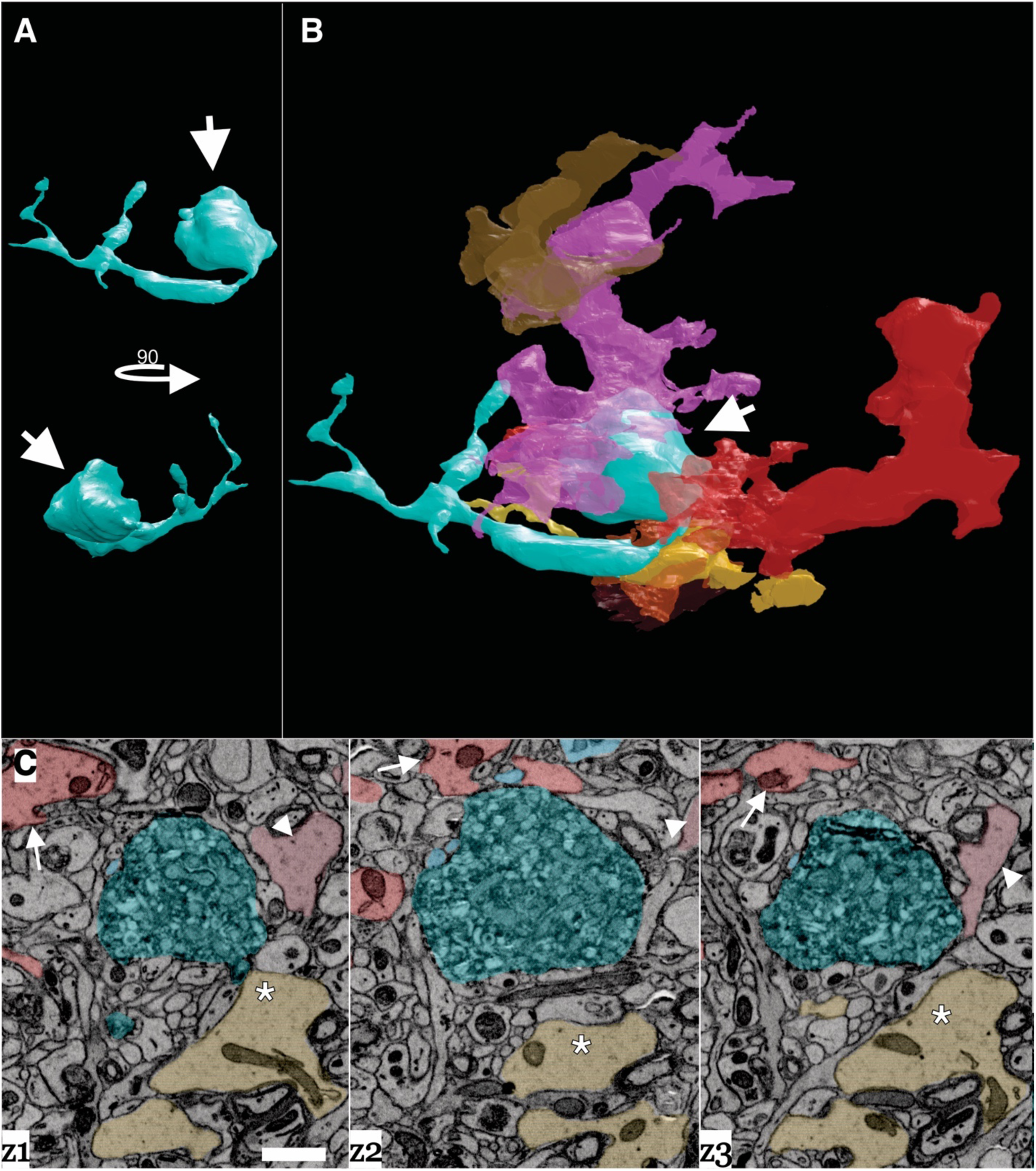
Dopamine axon swellings are surrounded by glia. (**A**) 3D reconstruction of a dopamine axon swelling from a cocaine sensitized mouse. Bottom image is rotated 90 degrees relative to top view. While arrows point to swelling. (**B**) 3D reconstruction of dopamine axon swelling (arrow) from (A) with glia surrounding it. Each differently colored object represents a different glial cell. (**C**) Montage of three serial EM images color coded to highlight the dopamine axon swelling (green), and three example glial cells (red/arrow, pink/arrow head, yellow/asterisk) that correspond to (A) and (B). Scale bar: (C) 1 µm

Finally, the tortuous and elongated nature of the mitochondria in these bulbs made us curious about possible mitochondrial changes in other cell types in the same tissue. One advantage of large volume EM datasets is that, since all cells and intracellular organelles like mitochondria are also labeled, we could ask whether cocaine altered mitochondrial length in other parts of DA neurons as well as other neurons in the NAc. Importantly, cyto-Apex2 staining did not occlude mitochondria (see Figure 1C, right panel for an example) thus allowing us to measure mitochondria lengths in cyto-Apex2 expressing DA axons.

We quantified mitochondrial lengths at five locations: in the NAc, we measured mitochondria length in: Apex2+ DA axons, MSN dendrites, and chemical synapse, likely glutamatergic axons (“CS axons”), and in the VTA, Apex2+ DA soma and Apex2+ DA dendrites (**Figure 9A**). Apex+ DA axons in the NAc had longer mitochondria throughout their arbors in the cocaine treated mice as compared to the saline controls. **Figure 9B** shows a representative 3D reconstruction of one such DA axon with its mitochondria from the cocaine (top and bottom left) and saline (middle, bottom right) treated mice. When quantified across numerous axons and across mice, we find that mitochondria in DA axons from cocaine treated mice are consistently longer than the saline controls (**Figure 9C**; mean ± SEM mitochondria length: +saline, 0.36 ± 0.01 µm, n = 162 mitochondria across 42 axons, 2 mice; +cocaine, 0.79 ± 0.05 µm, n = 162 mitochondria across 35 axons, 2 mice. P = 7.25e-25), suggesting that Apex2 expression alone does not cause mitochondrial elongation. When we extend this analysis to other neurons in the NAc, we found that cocaine also resulted in increased mitochondrial length in MSN dendrites, the putative targets of DA axons (**Figure 8D**; mean ± SEM mitochondria length (nm)/dendrite diameter (nm): +saline, 1.39 ± 0.12, n = 132 mitochondria across 50 dendrites, 2 mice; +cocaine, 3.0 ± 0.2, n = 260 mitochondria across 41 dendrites, 2 mice. P = 1.14e-6). However, increased mitochondria length appeared specific to DA axons and MSN dendrites, as we observed no difference in mitochondrial length in Apex-NAc CS axons between cocaine and saline treated mice (**Figure 8E**; mean ± SEM mitochondria length: +saline, 0.70 ± 0.05 µm, n = 104 mitochondria across 30 axons, 2 mice; +cocaine, 0.73 ± 0.04 µm, n = 164 mitochondria across 57 axons, 2 mice. P = 0.64). Surprisingly, we also did not observe any differences in mitochondrial length in VTA Apex+ DA dendrites or soma as compared to controls (**Figure 8F**; mean ± SEM mitochondrial length (nm)/dendrite diameter (nm) in Apex+ DA dendrites: +saline, 1.74 ± 0.25, n = 37 mitochondria across 4 dendrites, 1 mouse; +cocaine, 1.85 ± 0.22, n = 53 mitochondria across 10 dendrites, 1 mouse. P = 0.57; **Figure 8G**; mean ± SEM mitochondrial length in Apex+ DA soma: +saline, 2.46 ± 0.24 µm, n = 70 mitochondria across 4 soma, 1 mouse; +cocaine, 2.78 ± 0.23 µm, n = 141 mitochondria across 5 soma, 1 mouse. P = 0.68). Lastly, we confirmed that cocaine did not increase the frequency of mitochondria in DA axons (**Figure 9-Figure Supplement 1**; mean± SEM # mitochondria/µm: +saline, 0.14 ± 0.02, n = 96 mitochondria across 18 axons, 2 mice; + cocaine, 0.16 ± 0.02, n = 107 mitochondria across 20 axons, 2 mice. P = 0.71). Taken together, these results indicate that cocaine sensitization increases mitochondrial length in the mesolimbic pathway in a cell-type (i.e. in DA axons and not in CS axons) and sub-cellular specific (i.e. in DA axons and not DA soma or DA dendrites) manner.

**Figure 9.**
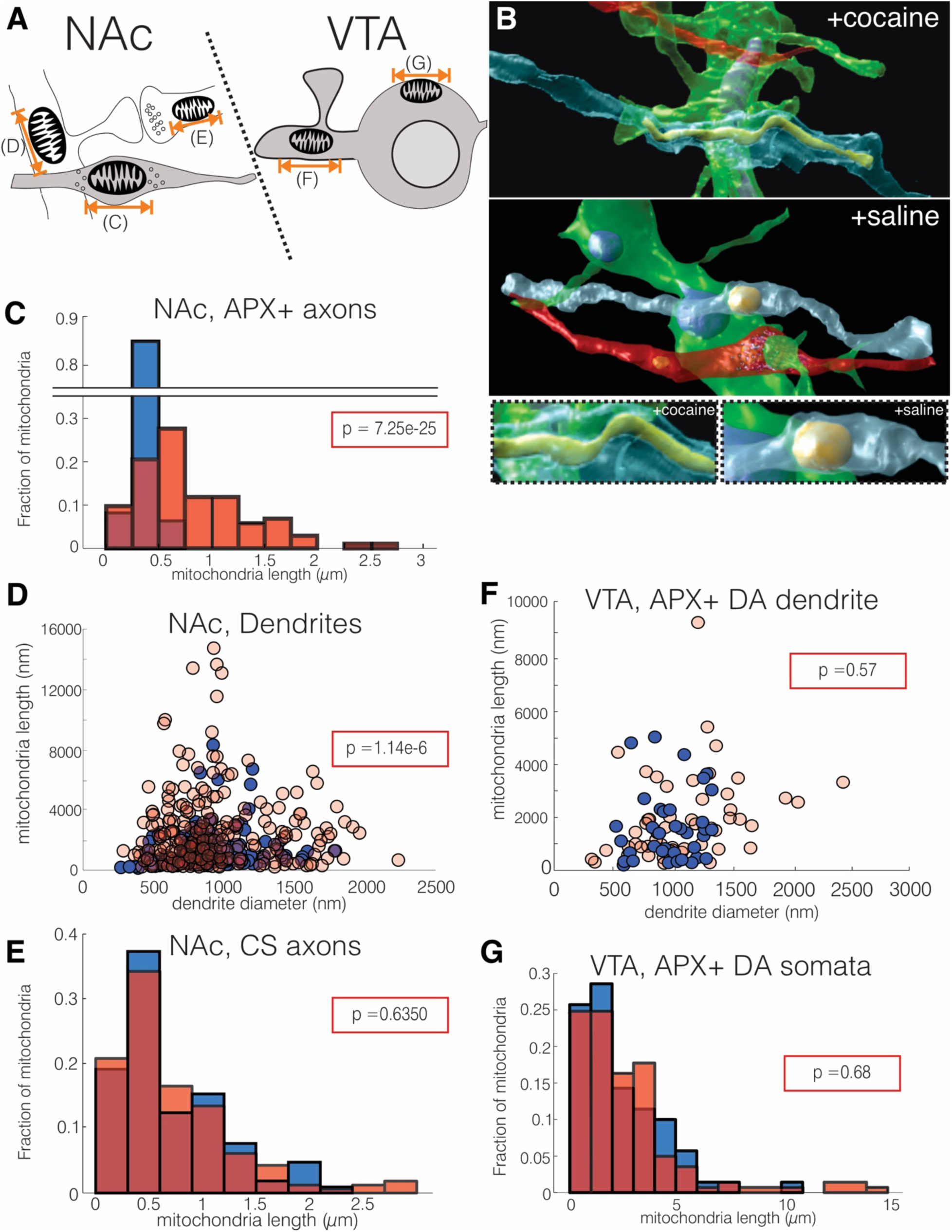
Cocaine results in increased mitochondria length in DA axons and MSN dendrites. (**A**) Cartoon depicting neurons where mitochondrial length was measured. *Left:* in the NAc, we measured mitochondria in: (C) Apex2+ DA axons, (D) MSN dendrites, and (E) excitatory axons making chemical synapses (CS axons). *Right*: in the VTA, we measured mitochondria in: (F) Apex2+ DA dendrites, and (G) Apex2+ DA soma. (**B**) 3D reconstructions from cocaine (top) and saline (bottom) treated mice of a DA axon (blue), afferent CS axon (red), and MSN dendrite (green) from the NAc. Shown is just the mitochondria reconstructed within the DA axon. Bottom images show zoomed views centered on the DA axon mitochondria from cocaine (left) and saline (right) treated mice. (**C**) histogram of mitochondria length in NAc, Apex2+ (“APX+”) DA axons (+saline: 0.36 ± 0.01 μm, n = 162 mitochondria across 42 axons, 2 mice; + cocaine: 0.79 ± 0.05 μm, n = 162 mitochondria across 35 axons, 2 mice. P = 7.25e-25. (**D**) Scatter plot of mitochondria lengths versus dendrite diameter of NAc MSN dendrites (+ saline: 1.39 ± 0.12 mito length (nm)/dendrite diameter (nm), n =132 mitochondria across 50 dendrites, 2 mice; + cocaine: 3.0 ± 0.2 μm mito length (nm)/dendrite diameter (nm), n = 260 mitochondria across 41 dendrites, 2 mice. P = 1.14e-6). (**E**) histogram of mitochondria lengths in NAc CS axons (+ saline: 0.70 ± 0.05 μm, n = 104 mitochondria across 30 axons, 2 mice; + cocaine: 0.73 ± 0.04 μm, n = 164 mitochondria across 57 axons, 2 mice. P = 0.64). (**F**) Scatter plot of mitochondria lengths versus dendrite diameter of VTA Apex2+ DA dendrites (+saline: 1.74 ± 0.25 mito length (nm) /dendrite diameter (nm), n = 37 mitochondria across 4 dendrites, 1 mouse; + cocaine: 1.85 ± 0.22 mito length (nm)/dendrite diameter (nm), n = 53 mitochondria across 10 dendrites, 1 mouse. P = 0.57. (**G**) Histogram of mitochondria length in Apex2+ DA Soma (+ saline: 2.46 ± 0.24 μm, n = 70 mitochondria across 4 soma, 1 mouse; + cocaine: 2.78 ± 0.23 μm, n = 141 mitochondria across 5 soma, 1 mouse. P = 0.68). Data: mean ± SEM. P-value: two-tailed Mann-Whitney U test.

## SUMMARY

We leverage the first large volume connectomic reconstruction of genetically labeled DA circuits to demonstrate that VTA DA neurons rarely make classical synapses in the medial shell of the NAc, resolving a long-standing question about whether DA communication occurs via specific synapses (Agnati *et al*., 1995; Bérubé-Carrière *et al*., 2012; Liu *et al*., 2018; Moss and Bolam, 2008; Omelchenko and Sesack, 2009; Rice and Cragg, 2008; Sesack et al., 1998). We find that most DA axons do not create classical synapses with targets in NAc and that DA axonal varicosities are primarily of four types, in descending order of frequency: 1. empty, 2. filled with small vesicles, 3. filled with large vesicles, or 4. filled with both small and large vesicles. Individual DA axons were more likely to contain stretches of a specific varicosity type than expected by chance, suggesting a previously unknown classification system for DA axons based on varicosity type. Additionally, the pleiomorphy of vesicles at individual varicosities suggests that DA axons may release different combinations of neurotransmitters at each varicosity as supported by evidence that some DA axons may release more than one neurotransmitter (Hnasko and Edwards, 2012; Sulzer et al., 1998; Sulzer and Rayport, 2000; Tritsch et al., 2016). At many DA varicosities, we instead found evidence of complicated three-dimensional membrane invaginations with nearby axons and dendrites – ‘spinules’ –previously reported as an alternative method of neuronal communication and plasticity at excitatory synapses in the hippocampus and cortex (see below). After a brief exposure to cocaine, we see widespread evidence of DA axonal rearrangements, including large scale axonal branching and the formation of blind ended ‘axonal bulbs’, filled with mitochondria, and surrounded by glia. Finally, we find longer mitochondria in some cell types (e.g. DA axons but not CS axons), and in some parts of neurons but not others (e.g. DA axons but not DA dendrites or soma).

### LIMITATIONS

Our results have multiple limitations that provide important context for interpreting our results. First, our cocaine sensitization protocol is a simplified version compared to more elaborate, behavioral protocols of drug addiction. While there are clear molecular and behavioral distinctions between cocaine sensitization and experimental models of addiction (Markou et al., 1999; McCutcheon et al., 2011), we chose cocaine sensitization as the minimum experimental model to first establish a baseline for the acute effects of cocaine. Follow-up studies that examine whether these alterations persist at longer time points or in models of addiction (e.g. self-administration) would shed further light on how these re-arrangements correlate with long term behavioral changes. Second, our genetic labeling approach targets all DA cell types indiscriminately within the VTA as defined by the efficiency of transgenic labeling or viral infection. We started these investigations as a necessary first step to evaluate expression, evaluate staining, and explore how connectomic datasets could be used to detail DA circuits. Future experiments where different DA cell types are differentially labeled (e.g., mitochondria and cytoplasmic-Apex2 targeted to two DA subtypes) in the same animal would further refine these initial results. Indeed, labeling of many potential DA cell types may have allowed us to further refine our classification system based on the composition of varicosities in individual axons (Figure 2-3). Finally, this study is limited in the number of animals used and the number of brain regions analyzed. DA axons are widely distributed in the brain and may have fundamentally different physical interactions with target neurons in other brain regions. Future investigations using the approach highlighted here in other brain regions could help address the generalizability of these results. While limited in numbers, however, this study represents one of the largest connectomic reconstructions to date (e.g. 3 control and 2 experimental animals) (Bates *et al*., 2020; Helmstaedter *et al*., 2013; Kasthuri et al., 2015b; Morgan et al., 2016; Morgan and Lichtman, 2020; Motta *et al*., 2019; Vishwanathan et al., 2017), and the differences we saw between control and cocaine exposed animals were large and consistent within cohorts across experimental conditions (i.e. no evidence of bulbs in DA axons in controls, and cocaine treated mice show extensive DA axon branching and specific mitochondrial changes).

## THE NANOSCALE ANATOMY OF DOPAMINERGIC CONNECTIONS

How neuromodulatory neurons interact with downstream targets remains largely unknown despite their large influence on brain function (Avery and Krichmar, 2017; Bargmann, 2012; Nadim and Bucher, 2014). By using Apex2 labeling and large volume automated serial EM, we have conducted the first ever large volume 3D nanoscale analysis of DA axon circuitry to thoroughly investigate the anatomy of DA axons. The existing literature characterizing DA axons has broadly relied on immunohistochemistry to label DA axons and varicosities in EM or fluorescence datasets. Previous EM studies have shown clear evidence for DA synapses on a variety of post-synaptic targets and locations, including dendritic spine heads, shafts, etc. (Nirenberg et al., 1997; Pickel et al., 1981; Uchigashima et al., 2016). However, many of these studies analyze smaller volumes of brains or often single EM sections, and therefore have not determined the frequency of classical ultrastructural synapses along individual DA axons; an important question when attempting to better understand DA circuitry and the relative contribution these identified synapses have to the broader circuitry of DA neurons. In addition, many of these studies were done in the rat, without genetic cell type labeling, and therefore examine boutons from DA neurons from mixed sources (e.g., substantia nigra in addition to VTA).

### VARICOSTIY DIVERSITY OF DOPAMINERGIC CONNECTIONS

In our analyses, we demonstrate that 62% of all DA axon varicosities quantified contain vesicles, but that very few (<2%) made any clear synapse with any neighboring neuron. Further, varicosities varied with the types of vesicles with a majority (38%) having no vesicles, 25% with small vesicles (∼ 48 ± 1.7 SEM nm in diameter), 19% with large vesicles (133 ± 5.5 SEM nm in diameter) and 18% with both. In all categories, DA axon varicosities contained far fewer vesicles than nearby excitatory boutons of comparable size (data not shown). As large core vesicles are associated with neuropeptide or high molecular weight neurotransmitters and small core vesicles with classical ionotropic neurotransmitters (Edwards, 1998), the presence of both types in DA axons, along with mixed size vesicle varicosities, suggests that DA axons, both as a population and at the level of individual axons, could release different neurotransmitters at different locations. In addition, we show that individual DA axons often contain multiple examples of a particular type of varicosity, more than expected by chance, suggesting axonal specificity in the types of potential transmission described above. Finally, fluorescent labeling of proteins involved in dopamine release (e.g. bassoon, TH, RIM1) showed that regions along DA axons were negative for dopamine but still positive for the molecular machinery necessary for dopamine release (Liu *et al*., 2018). Future experiments combining molecular details with serial EM (Fulton and Briggman, 2021; Liu and Cheng, 2016; Yamashita, 2016) could address whether these regions devoid of dopamine but positive for vesicle release machinery correlated with any of the varicosity types identified in this study (i.e. could locations with synaptic machinery but not dopamine correlate with Type I, empty varicosities described here).

### THE PAUCITY OF DA SYNAPSES

We find that only a small fraction (<2%) of all DA varicosities of any type (e.g. empty or not) made clear ultrastructural synapses with post-synaptic targets of any kind (i.e. spine, shaft, soma, etc.). The small number of DA boutons (6/410) with clear ultrastructural signs of a synapse were on the shaft and soma of resident NAc neurons, likely MSNs. While significant attention has been given to the spinous connections on MSNs, there is less data on dendritic shaft and somatic inputs onto these neurons, partly because dendritic spines are good proxies for synapses when using standard light microscopy approaches but there remain few good optical proxies for individual shaft or somatic synapses. A previous study demonstrated innervation of the shaft and soma of MSNs by boutons containing pleomorphic synaptic vesicles (Wilson and Groves, 1980), and we extend that work demonstrating that some of that innervation is likely from DA axons. We also found little evidence that DA varicosities of any type (e.g. empty or not) were more likely to be spatially proximate to spine synapses as suggested by previous studies as one possible model of targeted, non-synaptic DA transmission (Arbuthnott and Wickens, 2007; Sesack *et al*., 1998). Finally, a current theory of dopaminergic transmission is that while DA axons make specific ultrastructural synapses, there remains a ‘mismatch’ between presynaptic elements and post-synaptic dopamine receptors (Agnati *et al*., 1995), therefore invoking the idea of volume transmission. Our data clearly suggest differences with that model: the absence of ultrastructural evidence of DA synapses writ large. Future experiments combining, for example, either APEX tagged DA receptors or large volume immuno-electron microscopy (Fulton and Briggman, 2021) would address the proximity of different DA varicosity types to specific DA receptors.

### SPINULES AS A POSSIBLE MECHANISM OF DA TRANSMISSION

The other major type of direct physical interactions we found were numerous instances of axon and dendrites physically interdigitating with the membranes of DA varicosities. These interdigitations most closely resembled previously characterized synaptic structures termed “spinules” which occur in cortical and hippocampal neurons, and in both *in vitro* cell cultures and brain slices (Petralia *et al*., 2015; Spacek and Harris, 2004; Zaccard *et al*., 2020). Spinules are small (∼100 nm diameter) membranous protrusions typically found at or near synaptic sites at dendritic spine synapses. Spinules most often emerge from dendritic spines and to a lesser extent axonal boutons, but most often these spinules, regardless of their source, project into axons-either the presynaptic axon or a neighboring axon not actively forming a synapse (Spacek and Harris, 2004). Furthermore, spinules have been detected using high pressure freezing methods, free of any aldehydes, and in dissociated cultures of living neurons, suggesting that these invaginations are not artifacts caused by chemical fixation methods (Tao-Cheng *et al*., 2009; Zaccard *et al*., 2020). While their function is not totally known, spinule formation seems activity dependent (e.g. spinules are induced by K+ mediated neuronal depolarization, NMDA (Tao-Cheng *et al*., 2009), and glutamate activity (Richards et al., 2005)). Our results now show, for the first time, DA neurons also form spinules. However, unlike hippocampal neurons where spinules more often originate from dendrites, we find that 83% of spinules involving DA axons originate from unlabeled axons, presumably afferent axons from cortex or thalamus. Finally, we found that cocaine exposure does not change the frequency of spinules along DA axons (i.e. spinules/µm), but as DA axons branch more in response to cocaine, the newly formed axons also contain more spinules and thus more physical interactions with surrounding neurons. Labeling of other neuromodulatory pathways (e.g. cholinergic, serotonergic, etc.) could reveal whether such spinule likes structures are a common structural motif at neuromodulatory varicosities.

## COCAINE INDUCED DA CIRCUIT REMODELING

### Large scale DA axon remodeling

We find evidence of large-scale anatomical changes in DA axons following exposure to cocaine. Broadly, our data are consistent with active remodeling of dopamine axons: retraction bulbs on axon terminals in the process of pruning (i.e., removing connections) and increased axonal branching (i.e., formation of new connections). Additionally, coinciding with these structural plasticity events, we find evidence of alterations in mitochondria, potentially due to changes in cellular metabolism. While previous reports have focused on changes in DA spine density in the VTA and MSN dendritic spine synapse density in the NAc (Alcantara et al., 2011; Barrientos et al., 2018; Sarti et al., 2007), our results fill in an important gap to our understanding of how cocaine alters the structure of DA axons. Notably, these changes occurred after just 4 days of exposure and 4 days of withdrawal, suggesting a rapid pace of remodeling.

The most striking change in response to cocaine we discovered were large swellings in DA axons. These swellings bare striking similarities to remodeling events observed during development and traumatic brain injury (TBI). Axonal bulbs in development and TBI are distinguished from each other as either axon retraction or axon degeneration (*i.e.* “Wallerian degeneration”), respectively (Bishop *et al*., 2004; Johnson *et al*., 2013; Rosenthal and Taraskevich, 1977). We do not observe axon fragmentation nor removal of the entire axon, both hallmarks of axon degeneration, but rather the bulbs we observe are connected to otherwise intact DA axons and thus resemble more closely axonal pruning seen during development. These results raise several new questions: 1) what drives large bulb formation, 2) How are DA bulbs different from glutamatergic axon bulbs that are thought to form pruned synapses, given that DA axons rarely make conventional synapses, and 3) what, if any, release of neurotransmitter occurs at bulbs? Finally, we note that the largest difference in structural plasticity seen after exposure to cocaine were not changes at individual varicosities (i.e., more classical synapses per varicosity after cocaine exposure, more spinules after cocaine exposure, more or larger varicosities per length of axon after cocaine exposure). Instead, we find large scale changes in the arborization patterns of DA axons, like the plasticity observed over development. However, unlike developmental axonal arbor plasticity, which occurs generally as a monotonic decrease in the size of axonal arbors over development (Hubel et al., 1977; LeVay et al., 1980; Tapia et al., 2012), we see signs of both increased branching and possible retraction bulbs suggesting that different cellular mechanisms could mediate the plasticity described here. It would be interesting to see if other types of circuit plasticity follow similar principles.

### Glia and DA axonal remodeling

The presence of ‘activated’ glia around the DA axon bulbs further suggests these are areas of the DA axon being actively sensed and responded to by the brain’s glial monitoring system. While it is established that cocaine and other drugs of addiction can activate glial cells (Linker et al., 2019; Miguel-Hidalgo, 2009), the direct role for glia in mediating the effects of drug abuse has remained unclear. Here, for the first time, we provide direct anatomical evidence for a potential role for glia following cocaine exposure in mediating retraction bulb development and resolution, like how glial cells mediate axonal retraction bulbs during development (Bishop *et al*., 2004; Wilton *et al*., 2019).

### Cell type and cell compartment specific mitochondrial remodeling after cocaine exposure

In addition to observing signs of plasticity in DA axons, we also find evidence that cocaine can result in alterations at the subcellular level through changes in mitochondria. Previous studies have demonstrated that cocaine alters brain energy homeostasis, including changes in oxidative stress, cellular respiration, and enrichments of mitochondrial-related transcripts in NAc brain slice preps (Dietrich et al., 2005; Feng et al., 2014; Kalivas, 2009; Lehrmann et al., 2003; Volkow et al., 1991). The changes in mitochondrial length documented here could provide a potential cellular substrate for these energy changes (Glancy et al., 2020; Skulachev, 2001). Another possibility is that mitochondrial elongation is the result of hyperactivity of the mesolimbic circuit, as demonstrated by the increased movement of mice exposed to cocaine (Figure 6-Figure Supplement 1). Changes in mitochondrial cytochrome oxidase staining with increased neuronal activity are well known (i.e. cytochrome oxidase and ocular dominance columns in cortex) (Horton and Hedley-Whyte, 1984; Wong-Riley and Riley, 1983) and, if so, the cell-type specific nature of our results (i.e. DA axons and MSN dendrites show elongated mitochondria but excitatory, putative glutamate, axons and DA soma and dendrites of the VTA do not) suggest that cocaine may act preferentially on DA axons and MSNs, at least with regards to cellular metabolism. Finally, since many of these results could not have been collected in any other way, we conclude that large volume connectomic efforts with cell type specific labeling could be a valuable tool for the study of other neuromodulatory circuits.

## Materials and Methods

### Animals and AAV viruses

DAT-CRE mice ∼15 weeks used in this study were acquired from Xiaoxi Zhuang (The University of Chicago), and can also be found at Jackson Laboratory (https://www.jax.org/strain/020080). AAV-CAG-DIO-APEX2NES (Cyto-Apex) was acquired as a gift from the laboratory of Joshua Sanes (Harvard) and is now available at Addgene: #79907. AAV-CAG-DIO-APEX2-MITO was generated in our lab by cloning the mitochondrial targeting sequencing from mito-V5-APEX2 (Addgene #72480) and placing it on the 5’ end of APEX2-NES in AAV-CAG-DIO-APEX2NES. Finally, the nuclear export sequence (NES) was removed from APEX2. AAV9 virus was generated at the University of North Carolina School of Medicine Vector Core facility (https://www.med.unc.edu/genetherapy/vectorcore/). Animal care, perfusion procedures, and AAV injections were followed according to animal regulations at the University of Chicago’s Animal Resources Center (ARC) and approved IACUC protocols.

### AAV injections

AAV9 injections were performed using a standard stereotaxic frame. 70-100 nl of virus (∼2.9×10^12^ viral genomes/ml) were bilaterally injected into the VTA using the stereotactic coordinates: 3.1 posterior of bregma, 0.55 lateral bregma, and 4.4 ventral of the dura. Mice were aged 4 weeks, based on prior experience with APEX expression (Sampathkumar *et al*., 2021) to allow for AAV expression before perfusion or cocaine sensitization experiments.

### Cocaine sensitization

Mice were given a once daily intraperitoneal (IP) injection of either cocaine (10 mg/kg) or equivalent volume of saline every other day for a total of 4 injections. Immediately following each injection, mice were place in a novel environment where their locomotor activity was automatically monitored for one hour and then returned to their home cage.

### Apex2 staining and EM preparation

Brains were prepared in the same manner and as previously described (Hua *et al*., 2015). Briefly, an anesthetized animal was first transcardially perfused with 10ml 0.1 M Sodium Cacodylate (cacodylate) buffer, pH 7.4 (Electron microscopy sciences (EMS) followed by 20 ml of fixative containing 2% paraformaldehyde (EMS), 2.5% glutaraldehyde (EMS) in 0.1 M Sodium Cacodylate (cacodylate) buffer, pH 7.4 (EMS). The brain was removed and placed in fixative for at least 24 hours at 4°C. To ensure that the same brain region was isolated across replicates, the brain was mounted to the vibratome on the anterior side, and carefully trimmed down through the cerebellum until it just touched the posterior cortex. The brain was then completely sectioned into a series of 300 µm vibratome sections, placed, in order, into a 24 well dish, and incubated into fixative for 24 hours at 4°C. Apex2 precipitation and polymerization was performed by washing slices extensively in cacodylate buffer at room temperature, incubated in 50mg/ml 3,3’diaminobenzidine (DAB) for 1 hour at room temperature followed with DAB/0.03% (v/v) H_2_0_2_ until a visible precipitate forms (15-20 minutes). Slices were washed extensively in cacodylate buffer. Slices were reduced in 0.8 %(w/v) Sodium Hydrosulfite in 60%(v/v) 0.1 M Sodium Bicarbonate 40 %(v/v) 0.1 M Sodium Carbonate buffer for 20 minutes and washed in cacodylate buffer. To ensure the same region of the Nucleus Accumbens was taken across animal samples, we used areal landmarks of the anterior commissure, shape of the corpus callosum, and boundary of positive and negative Apex2 staining in the medial portion of the brain to isolate the medial shell of the NAc. The medial shell of the NAc and a piece of tissue spanning the entire VTA were excised and prepared for EM by staining sequentially with 2% osmium tetroxide (EMS) in cacodylate buffer, 2.5% potassium ferrocyanide (Sigma-Aldrich), thiocarbohydrazide, unbuffered 2% osmium tetroxide, 1% uranyl acetate, and 0.66% Aspartic acid buffered Lead (II) Nitrate with extensive rinses between each step with the exception of potassium ferrocyanide. The sections were then dehydrated in ethanol and propylene oxide and infiltrated with 812 Epon resin (EMS, Mixture: 49% Embed 812, 28% DDSA, 21% NMA, and 2.0% DMP 30). The resin-infiltrated tissue was cured at 60°C for 3 days. Using a commercial ultramicrotome (Powertome, RMC), the cured block was trimmed to a ∼1.0mm x 1.5 mm rectangle and ∼2,000, 40nm thick sections were collected from each block on polyamide tape (Kapton) using an automated tape collecting device (ATUM, RMC) and assembled on silicon wafers as previously described (Kasthuri *et al*., 2015a). The serial sections were acquired using backscattered electron detection with a Gemini 300 scanning electron microscope (Carl Zeiss), equipped with ATLAS software for automated wafer imaging. To further ensure we captured the same region of the NAc across datasets, we used the anterior commissure, which was contained within our serial sections, as a guide for were to reproducibly place the ROI during imaging. Dwell times for all datasets were 1.0 microsecond. For 20nm and 6nm resolution data sets, sections were brightness/contrast normalized and rigidly aligned using TrakEM2 (FIJI) (Cardona et al., 2012) followed by non-linear affine alignment using AlignTK (https://mmbios.pitt.edu/aligntk-home) on Argonne National Laboratory’s super computer, Cooley. Different image processing tools were packaged into Python scripts that can be found here: https://github.com/Hanyu-Li/klab_utils. Final volumes collected for each mouse were: 1. Mito-Apex = 160 µm x 160 µm x 11 µm @ 6nm, 1 mouse, 2. Cyto-Apex = 450 µm x 350 µm x 30 µm @ 20nm, 4 mice and 130 µm x 130 µm x 10 µm @ 6nm, 4 mice.

### Data Analysis

Aligned datasets were manually skeletonized and annotated using the publicly available software, Knossos (https://knossos.app). All quantifications reported are mean ± standard error of the mean (SEM). Two-tailed Mann Whitney U statistics test was used to test for significance (Marx et al., 2016). Data annotations were done by two individuals (GAW, AMS). To ensure accuracy of the data, ∼33% of the annotations from one person was verified by the other. We found a >98% agreement between manual annotators. Classes of cell types were identified by distinguishing anatomical properties: Apex2+ DA neurons (Soma and dendrites in the VTA and axons in NAc) were identified by their dark precipitate, Medium spiny neurons in the NAc by the presence of dendritic spines, and chemical synapse axons in the NAc by their formation of synapses on dendritic spines. Skeleton information was exported into comma separated matrices using homemade Python scripts that compute skeleton features from the Knossos xml annotation file. Code is freely available here: https://github.com/knorwood0/MNRVA. Quantification and plotting of different anatomical features were performed in Matlab and excel.

### DA axon varicosities

DA axon varicosities were identified as regions where: 1. the DA axon diameter increased ∼3 fold (∼100nm to ∼300 nm in axon diameter) in the absence of a mitochondria, and 2. regions with increased diameter that contained both a mitochondria and cloud of vesicles. Areas of axon enlargement containing just a mitochondrion were not marked as a varicosity. Small vesicles were scored as those ranging in diameter of 26-67 nm and large vesicles ranged in diameter of 90-238 nm.

### DA axon spinules

Spinules in the mito-Apex2 datasets were identified as regions of membrane invaginations into DA axons that could be traced out to a nearby neurite. Spinules in cyto-Apex2 datasets were identified as invaginations that were void of Apex2 staining that invaginated into the Apex2+ DA axon and could be traced out to a nearby neurite. Spinule invaginations were further confirmed by observing a neurite entering a DA axon and disappearing in the image stack using the three orthogonal views of Knossos. Neurites that passed over DA axons were not scored as contact points/spinules.

### Monte Carlo Simulations of DA axon varicosity types

Monte Carlo simulations were ran using custom scripts in Matlab and are freely available upon request.

### DA axon branching

Cyto-Apex2 DA axons were manually reconstructed and the number of branches each axon made was scored.

### Large DA axon swellings

The entire volume of each dataset (∼0.5 x 0.5 x 0.0 2 mm) was surveyed for the presence of retraction bulbs. Additionally, the entire EM section (∼1×1.5mm) was spot checked by manually looking across every 20^th^ section.

### Glia identification

Glia were identified by several criteria: extensive branching, cytoplasmic granules, vacuoles, and lack of any synapses.

### Mitochondria length

The length of mitochondria was measured in Knossos by measuring along the longest axis from one of the three orthogonal views (xy,xz,yx). Dendrite diameter was measured by centering the mitochondria in the Knossos viewing window and then taking the average diameter between all three orthogonal axes.

### 3D rendering

3D manual segmentation was done using VAST (Berger et al., 2018) and exported using MATLAB as .OBJ files which were then imported and rendered using Blender 2.79.

### Data availability

All EM data and reconstructions are freely available upon request. We currently store the data on our local servers and will offer it to be hosted on public databases including https://neurodata.io/ and NeuroMorpho.org.

---

**Figure 3-Figure Supplement 1.**
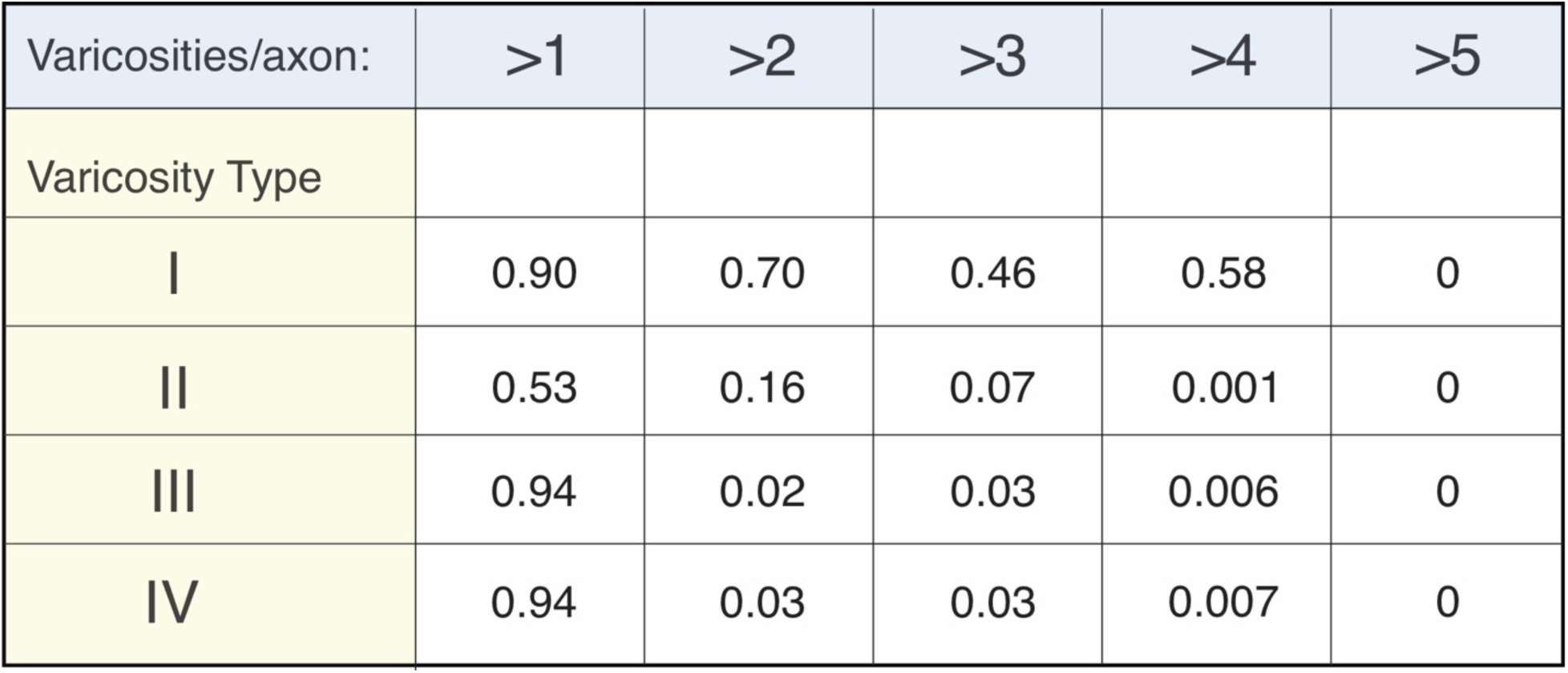
Table of P-values for Monte Carlo simulations. The top row (light blue) represents each data point in the x-axis of Figure 3B that counts the number of axons containing greater than the specified number of varicosities/axon. The left column (light yellow) separates each varicosity type. P-values were calculated by dividing the number of times the simulated axon had a specified varicosity type (yellow column) appear more than the number of specified times (blue row) by the total number of monte carlo simulations (100,000).

**Figure 5 - Figure Supplement 1.**
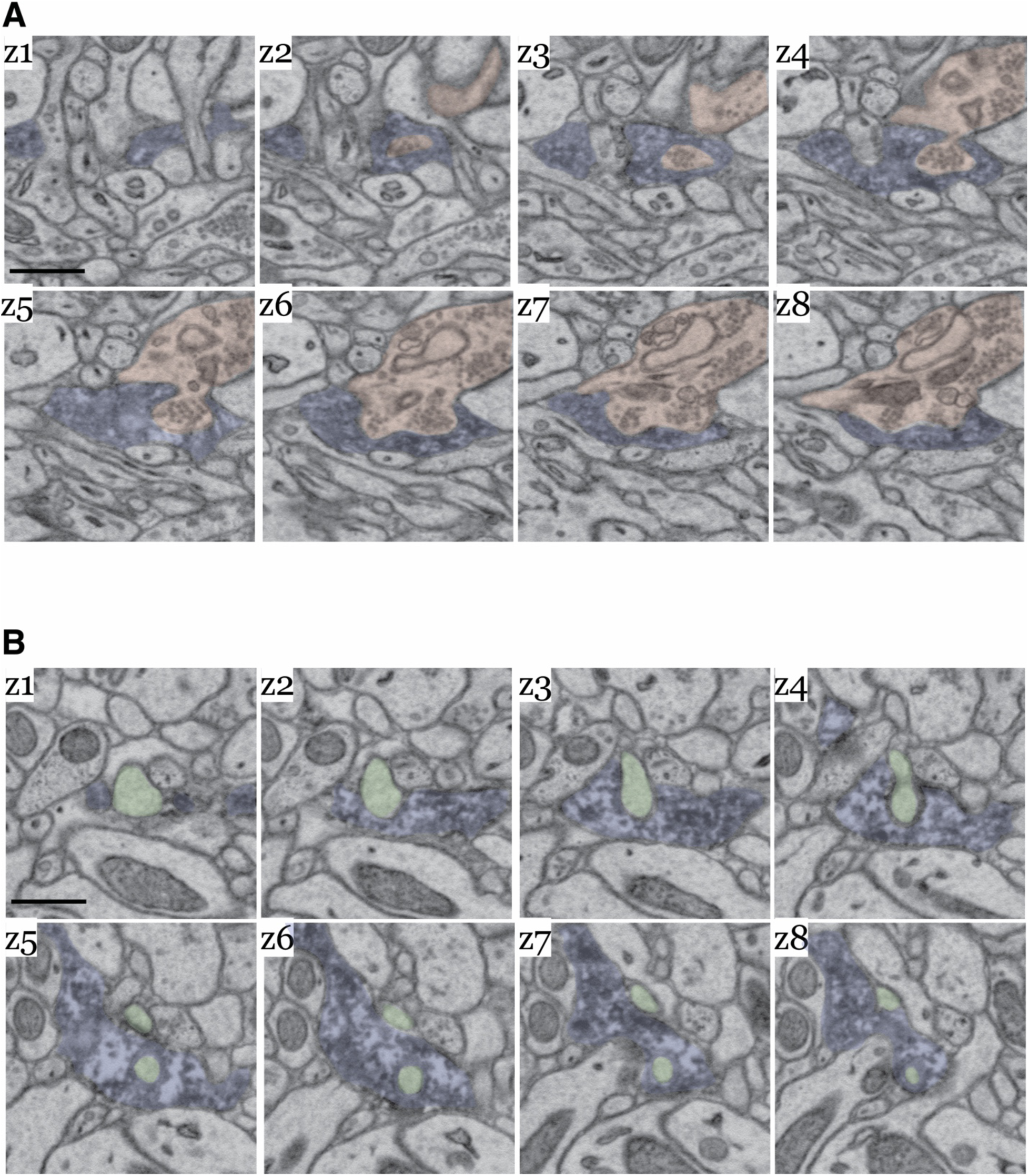
Image montage of DA axon interdigitations. (**A**) EM z-series of the interdigitation between a DA axon (blue) and vesicle-filled CS axon (red) that corresponds to the 3D rendering in Figure 5B, left image. (**B**) EM z-series of the interdigitation between a DA axon (blue) and NAc dendritic spine (green) that corresponds to the 3D rendering in Figure 5B, right image. Scale bar = (A,B) 1 µm.

**Figure 6-Figure Supplement 1.**
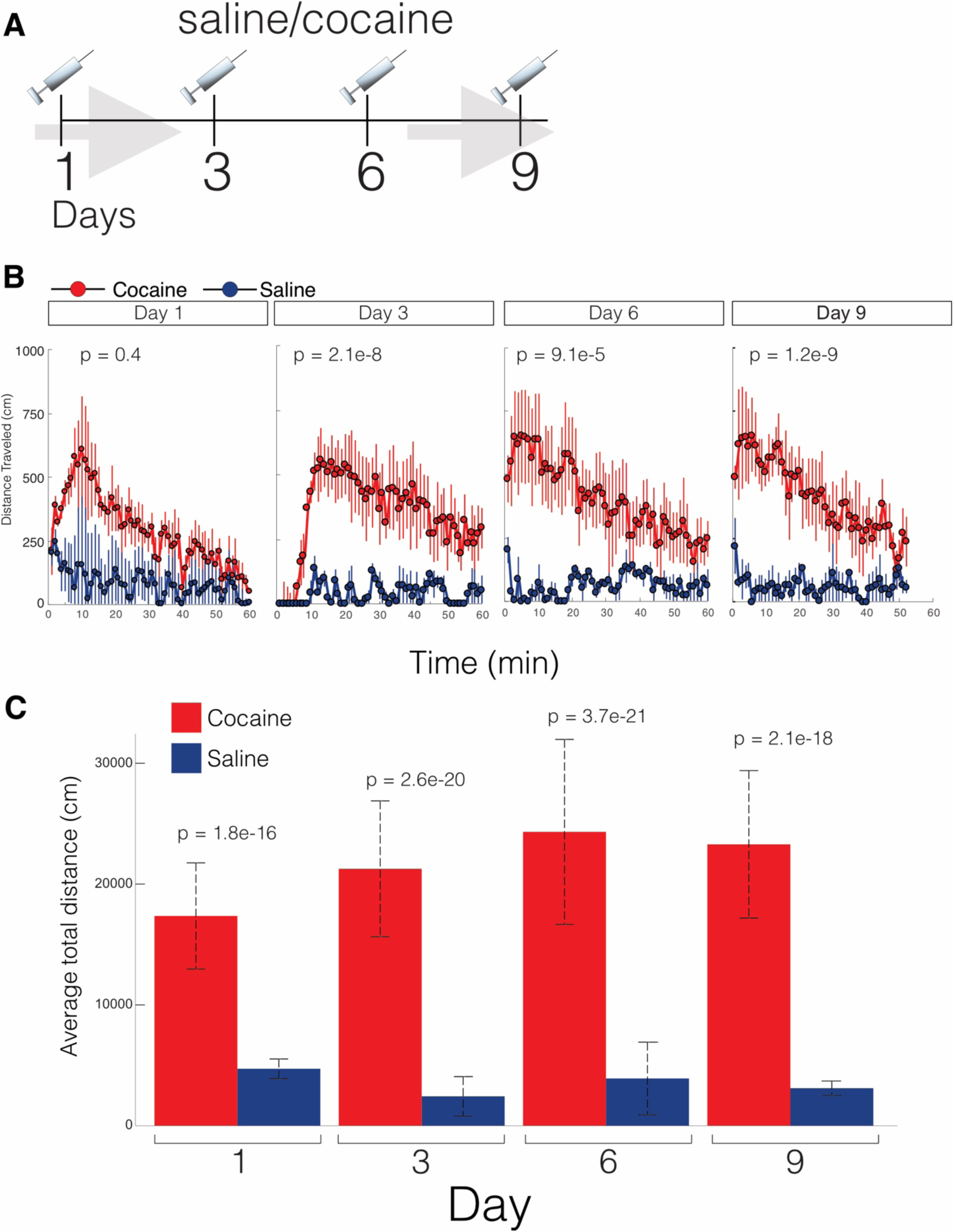
Cocaine treated mice have increased locomotor activity. (**A**) Cartoon depicting cocaine sensitization protocol: Mice received a single intraperitoneal (IP) injection of cocaine (10 mg/kg) or equivalent volume of saline, placed in a novel cage environment, and their locomotor activity was monitored for 1 hr. This procedure was repeated every 3rd day for 4 rounds of injections. 4 days following the last injection (e.g. Day 13), mice were perfused and brains processed for Apex2/EM staining (see Methods). (**B**) Scatter plot of the average distance traveled (cm) versus time for the first day of cocaine or saline injection. (**C**) Total average velocity (cm) of mice immediately following IP cocaine or saline injections. For cocaine and saline, n = 3 mice each; data = mean ± SEM. P-value: two-tailed Mann-Whitney U test.

**Figure 6-Figure Supplement 2.**
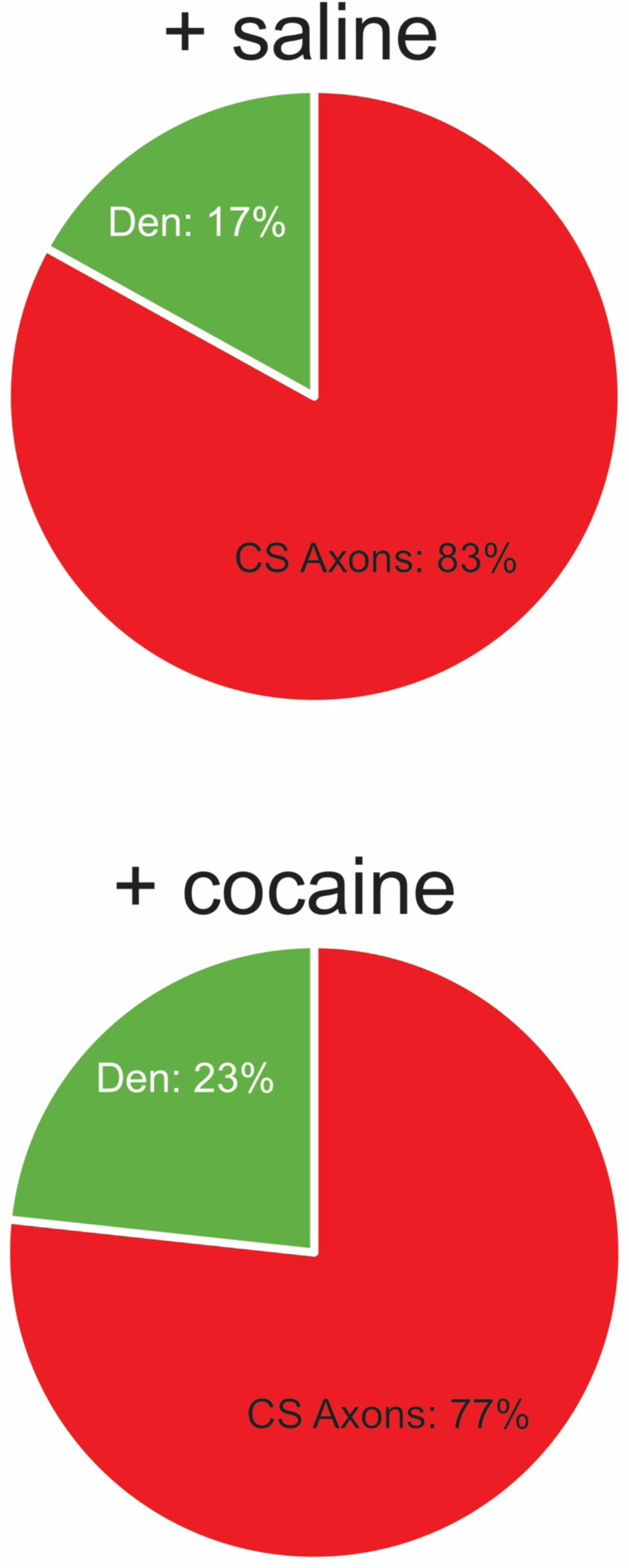
Cocaine does not change the proportion of Dopamine targets. Pie charts showing proportion of contact points (i.e. “spinules”) that are with CS axons (red) and dendrites (green) in saline (top) and cocaine (bottom) treated mice (+ saline: 83% (49/59) axo-axonic, 17% (10/59) axo-dendritic, n= 19 DA axons with 59 contact points scored; + cocaine: 77% (56/73) axo-axonic, 23% (17/73) axo-dendritic, n = 20 DA axons with 73 contact points scored).

**Figure 9-Figure Supplement 1.**
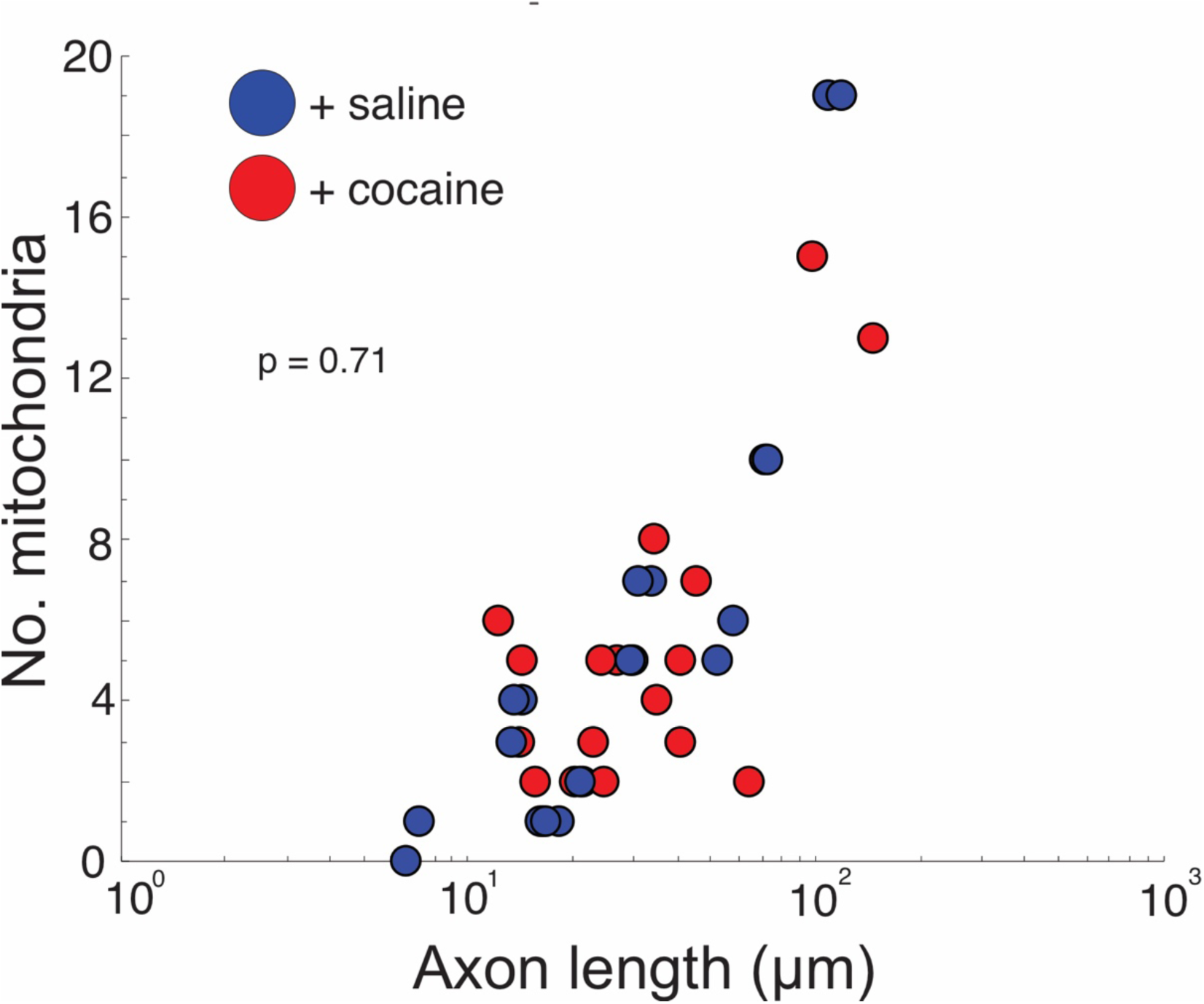
Cocaine does not change the number of mitochondria in DA axons. Scatter plot of the number of mitochondria versus length of axon (μm). (+ saline: 0.14 ± 0.02 mitochondria/µm length of axon, n = 96 mitochondria counted across 18 axons, 2 mice; + cocaine: 0.16 ± 0.02 mitochondria/µm length of axon, n = 107 mitochondria counted across 20 axons, 2 mice. P = 0.71). Data: mean ± SEM. *P*-values: two-tailed Mann-Whitney U test.

